# Proximity labelling of the BAK macropore uncovers a new role for SLC35A4-MP in mitochondrial dynamics

**DOI:** 10.64898/2026.03.22.713508

**Authors:** Matthew P. Challis, Stephanie M. Molé, Saveen Giri, Ryker Dumbrill, Matthew J. Eramo, Alice J. Sharpe, Sarah E. J. Morf, Kate McArthur, Luke E. Formosa, Michael T. Ryan

**Author notes:** The University of Sydney, New South Wales 2006, Australia (Michael T. Ryan). Co-corresponding authors: Kate McArthur, Luke E. Formosa and Michael T. Ryan.

## Abstract

Mitochondrial permeabilization by the apoptotic executioners BAK and BAX represents a critical stage of mitochondrial apoptosis and facilitates the release of pro-inflammatory mitochondrial DNA via herniation of the inner mitochondrial membrane. This study utilises TurboID proximity labelling to investigate the temporal changes of the BAK proximal proteome during mitochondrial herniation and apoptosis. In doing so, we detail a comprehensive BAK proximal proteome, both at steady state and during apoptosis and observe the loss of MICOS complex stability and proximity to the BAK pore as apoptosis proceeds. In addition, we identify the mitochondrial microprotein SLC35A4-MP proximal to the BAK pore and reveal a SLC35A4-MP dependent modulation of OPA1 processing. Furthermore, loss of SLC35A4-MP delays mitochondrial fragmentation in response to a variety of stressors, uncovering a previously unrecognised role for SLC35A4-MP in fine-tuning mitochondrial rearrangement during apoptotic stress.

## Introduction

Mitochondria are central to the execution of key signalling pathways including programmed cell death. Activation and oligomerisation of the pore forming pro-apoptotic BCL2-family proteins, BAK and BAX, represents a critical checkpoint for intrinsic apoptosis. BAK/BAX oligomerisation at the mitochondrial (OMM) causes mitochondrial outer membrane permeabilization (MOMP) and ultimately facilitates the release of pro-apoptotic factors, including cytochrome *c* into the cytosol ^(1)^. Once in the cytosol, cytochrome *c* engages the apoptotic protease-activating factor, Apaf-1, to form the apoptosome which cleaves and activates the initiator caspase 9, activating the executioner caspases 3 and 7 to accelerate cell death processes ^(2)^. In tandem to this, formation of BAK/BAX macropores in the outer mitochondrial membrane (OMM) leads to “herniation” of the inner mitochondrial membrane (IMM) into the cytosol ^(3–5)^. IMM herniation facilitates the release of the mitochondrial genome (mitochondrial DNA; mtDNA) into the cytosol, where it acts as a Danger Associated Molecular Pattern (DAMP) and engages cytosolic DNA sensors ^(3, 5)^. Ordinarily, apoptotic caspases restrain the immunogenic capacity of cytosolic mtDNA by inactivating cytosolic immune effectors and keeping apoptosis immunologically silent ^(6)^. However, in instances where pharmacological inhibition, or sub-lethal levels of MOMP (termed minority MOMP) render caspases insufficient to execute cell death ^(7)^, cytosolic mtDNA promotes a Type I Interferon (IFN) response, resulting in the production of pro-inflammatory cytokines ^(6, 8, 9)^. MtDNA associated inflammation has been implicated in numerous disorders, including Systemic Lupus Erythematosus ^(10)^ and models of Parkinson’s disease ^(11, 12)^, highlighting the importance of understanding the mechanisms by which mtDNA can escape.

IMM herniation and subsequent mtDNA release is facilitated by a dramatic remodelling of the highly folded mitochondrial cristae. Significant bending of cristae membranes occurs at two major sites: the terminal cristae tips – mediated by groups of F_1_F_0_ -ATPase dimers ^(13)^ – and the intersection of the cristae membrane and inner boundary membrane, known as the cristae junction ^(14)^. Cristae junctions are stabilised by a multi-subunit protein complex known as the Mitochondrial Contact Site and Cristae Organising System (MICOS) that connects with the sorting and assembly machinery (SAM) complex in the OMM to form the mitochondrial intermembrane space bridging complex (MIB) ^(15–18)^. In addition, oligomers of the large inner membrane GTPase Optic Atrophy 1 (OPA1) have been shown to be important for maintaining cristae architecture, particularly in regulating cristae width ^(19)^. There are several OPA1 isoforms, including membrane anchored long OPA1 (L-OPA1) isoforms, and proteolytically cleaved soluble short OPA1 (S-OPA1) isoforms that together, regulate IMM fusion and maintenance of the cristae ^(20, 21)^. Both OPA1 and the MICOS complex are targets for the mitochondrial response to stress. Stress conditions activate proteolytic conversion of L-OPA1 into S-OPA1, blocking IMM fusion and causing fragmentation of the mitochondrial network ^(22)^. In addition, stress results in cleavage of the MICOS subunit MIC19, leading to MICOS-MIB disassembly ^(23)^. How the IMM, and its integral protein complexes are reorganised during a stress such as IMM herniation is not clear, nor is it understood whether any integral IMM proteins are required for regulating morphological changes to the IMM during herniation.

To identify the repertoire of proteins moving in and out of proximity with the BAK pore during mitochondrial herniation, we re-expressed full length mouse BAK fused to the fast acting, promiscuous biotin ligase TurboID ^(24)^ in *Mcl1^-/-^Bak^-/-^Bax^-/-^*mouse embryonic fibroblasts (MEFs). This approach explored the spatial dynamics of mitochondrial proteins during mitochondrial herniation and identified a subset of IMM proteins that move into proximity with the BAK macropores. Subsequent characterisation of a 103 amino acid microprotein, SLC35A4-MP, demonstrated modulation of OPA1 processing and defects in mitochondrial fragmentation – highlighting the utility of this approach, and providing insight into the fine-tuning of mitochondrial dynamics during apoptotic stress.

## Results

To investigate the dynamic process of mitochondrial herniation, we utilised our HA epitope-tagged TurboID-BAK fusion protein (TurboBAK), stably expressed in *Mcl1^-/-^Bak^-/-^Bax^-/-^*MEFs to identify proteins proximal to BAK. The fast activity of the TurboID biotin ligase (biotin labelling within 10 min) allows us to sample the BAK proximal proteome at different timepoints following BAK activation. BAK pore formation was induced by treating cells with the pro-apoptotic BH3 mimetic ABT-737, in the absence of the pro-survival protein MCL1. The addition of pan-caspase inhibitor QVD-OPh prevents the caspase mediated progression of apoptosis, allowing for herniation to proceed (Fig. 1A) ^(3)^. We began by assessing the functionality of TurboBAK in comparison to our control *Mcl1^-/-^* MEFs. When assessed by Blue-Native (BN)-PAGE (Fig. 1B), TurboBAK assembled into a complex consistent with the 440 kDa inactive BAK complex ^(25)^ and following treatment with ABT-737, TurboBAK disassembled from the inactive complex (Fig. 1B) with the concomitant activation of caspases 3 and 7 (Fig. 1C). Whilst TurboBAK was able to initiate apoptosis, activation was slower than endogenous BAK in the *Mcl1^-/-^* control MEFs (Fig. 1B-C). Despite the slower kinetics, hallmarks of apoptosis including TurboBAK clusters, mitochondrial network collapse and herniation of mtDNA through condensed TOM20-positive cup-like structures was observed by 3 h post apoptosis treatment (Fig. 1D inset)^(3, 26)^. Even though the apoptotic kinetics of TurboBAK-expressing cells were delayed compared to *Mcl-1^-/-^* control cells, which expressed both endogenous BAK and BAX, we reasoned that the 1 – 2 h delay was advantageous, allowing for a clearer delineation between apoptotic initiation, BAK pore formation and mitochondrial herniation in the TurboBAK-expressing cell line.

**Figure 1:**
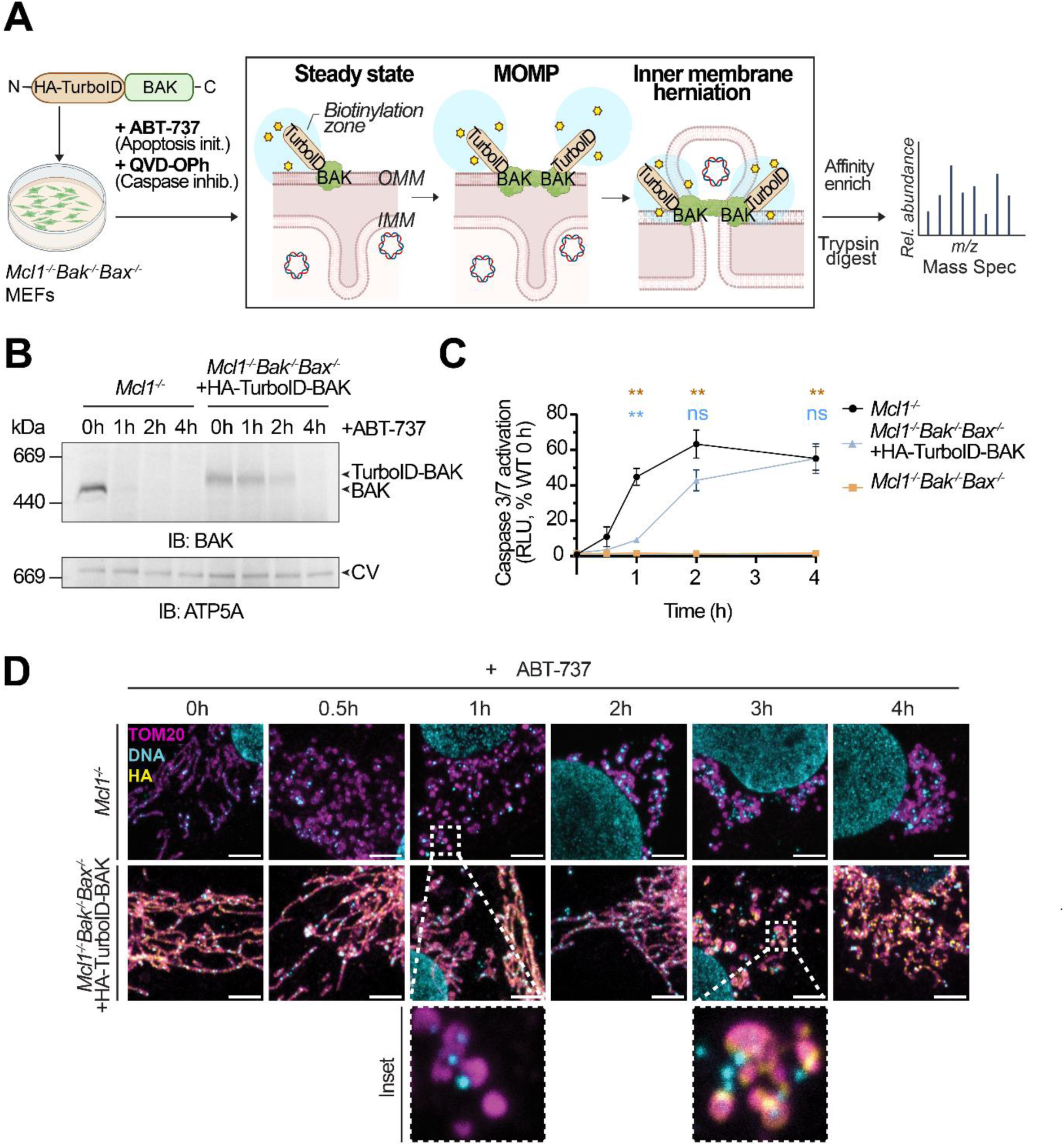
TurboBAK recapitulates key apoptotic events with delayed kinetics. **A.** Schematic illustrating the strategy used to investigate the proteomic environment of the BAK pore during apoptosis and mitochondrial herniation. **B.** The native, inactive BAK complex disassembles following activation by ABT-737 [2 µM] in control *Mcl1^-/-^* and *Mcl1^-/-^Bak^-/-^Bax^/-^*MEFs expressing HA-TurboID-BAK. Whole cells were solubilised with 1% digitonin and native BAK complexes were resolved by BN-PAGE and Western blot analysis. Complex V (ATP5A) was used as a loading control. **C.** Temporal analysis of caspase 3/7 activity in *Mcl1^-/-^, Mcl1^-/-^Bak^-/-^Bax^-/-^*, or *Mcl1^-/-^Bak^-/-^Bax^-/-^* + HA-TurboID-BAK MEFs treated with ABT-737 [2µM], relative to the 0 h *Mcl1^-/-^* control. Data points indicate mean ± SEM from n = 4 independent experiments. * = *p* < 0.05, ** = *p <* 0.01 by 2-way ANOVA with Tukey’s multiple comparison test, compared to the *Mcl1^-/-^* control. **D.** Representative confocal images of *Mcl1^-/-^Bak^-/-^Bax^-/-^* + HA-TurboID-BAK MEFs following treatment with ABT-737 [2 µM] and QVD-OPh [20 µM]. Immunofluorescence was performed using anti-TOM20 (magenta, OMM) and anti-DNA (cyan) antibodies to visualise mtDNA extrusion events. Anti-HA (yellow) was used to visualise HA-TurboID-BAK localisation. Scale bars indicate 4 µm.

Next, we used mass spectrometry to identify biotinylated proximal proteins labelled by TurboBAK at different timepoints throughout mitochondrial herniation (0 h steady state, 1 h pre-MOMP, 2 h post-MOMP and 4 h post herniation). The number of enriched proteins was consistent at each timepoint, although the number of enriched mitochondrial proteins was reduced at later time points (Fig. 2A). Initially, we sought to characterise the BAK proximal proteome at steady state (0 h, in the absence of ABT-737). Gene Ontology (GO) terms associated with mitochondrial membrane organisation, fission, localisation and transport were all enriched, consistent with specific labelling of the OMM (Fig. 2B). Furthermore, network analysis of the steady state BAK proximal proteome, revealed proteins associated with protein ubiquitination, peroxisome biogenesis, proteasome and protein transport in addition to the mitochondria specific complexes (Fig. S1A).

**Figure 2:**
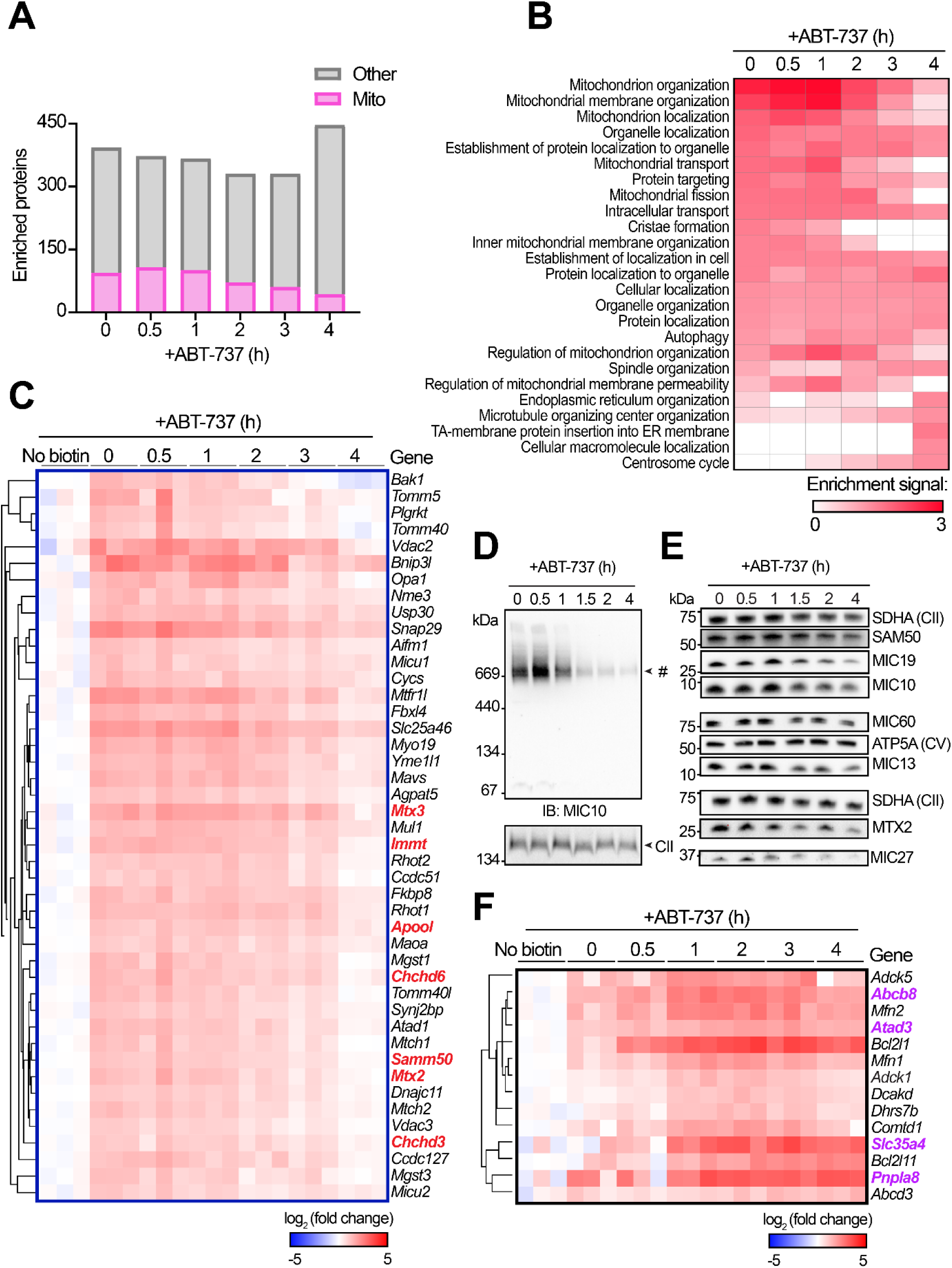
Mitochondrial herniation leads to decreased stability of the MICOS complex and enrichment of specific inner membrane proteins at the BAK pore. **A.** Number of enriched proteins in *Mcl1^-/-^Bak^-/-^Bax^-/-^* cells expressing TurboBAK at each time point post treatment, relative to no biotin control. Proteins with a log_2_ (fold change) > 1, and a *p* value < 0.05 were considered significantly enriched. **B.** Top enriched GO: Biological process terms at each time point of the apoptotic time course. **C.** Cluster of mitochondrial proteins with decreased enrichment over the duration of apoptosis, indicating reduced proximity to TurboBAK. Components of the MICOS-MIB are highlighted in red. **D.** Blue-Native Page and immunoblot analysis to observe MICOS complex assembly (IB: MIC10) from isolated mitochondria solubilised in 1 % digitonin following initiation of caspase inhibited apoptosis. # = MICOS complex. Complex II (SDHA) was used as a loading control. Representative of n = 2 independent experiments **E.** SDS-PAGE and immunoblot analysis of individual MICOS-MIB components from isolated mitochondria following treatment with ABT-737 and QVD-OPh. SDHA and Complex V ATP5A serve as loading controls. Representative of n = 2 independent experiments. **F**. Cluster of mitochondrial proteins that have increased enrichment over the apoptotic time course, indicating enhanced proximity to TurboBAK. Proteins localised to the IMM are highlighted in purple.

After 4 h ABT-737 treatment, the enrichment of GO terms associated with mitochondrial membrane organisation, localisation and fission were all lost. Conversely, the enrichment of non-mitochondrial terms associated with endoplasmic reticulum (ER) and microtubule organisation increased over time (Fig. 2B), consistent with their recruitment to mitochondria to facilitate mitochondrial fragmentation and peri-nuclear clustering during apoptosis ^(27, 28)^. In addition, this proximity labelling approach allowed us to capture dynamic changes in the enrichment of proteins associated with autophagy and the regulation of mitochondrial membrane permeability. Upon closer examination of the contributing proteins, we observed enrichment of the cytosolic autophagy adaptors OPTN1 and SQSTM1 (Fig. S2A), along with BCL-2 family proteins *Bcl2l1* (BCL-XL) and *Bcl2l11* (BIM) (Fig. S2B), consistent with previous reports detailing their dynamic recruitment to, or interaction with the BAK pore during apoptosis and mitochondrial herniation ^(26, 29, 30)^.

Given our interest in how the mitochondria is remodelled during MOMP and IMM herniation, and the observed changes in enrichment of mitochondrial GO terms, we focused our analysis on the mitochondrial proteins significantly enriched over the course of BAK activation. Hierarchical clustering of all significantly enriched mitochondrial proteins (at any timepoint) revealed distinct clusters of proteins that had changes in enrichment during herniation, indicative of a change in proximity to the BAK pore (Fig. S2C). One such cluster was enriched in proteins of the MICOS-MIB complex, with enrichment of MICOS subunits MIC60 (*Immt*), MIC27 (*Apool*), MIC25 (*Chchd6*), MIC19 (*Chchd3*) and SAM complex subunits SAM50 (*Samm50)*, MTX2 (*Mtx2)* and MTX3 (*Mtx3)* declining over time following BAK activation (Fig. 2C & S2C; blue box). BN-PAGE analysis of MIC10 assembly into the MICOS complex in control *Mcl1^-/-^* MEFs revealed a depletion of the fully assembled MICOS complex after 1.5 h (Fig. 2D), suggesting that MICOS stability is compromised following apoptotic initiation. Immunoblotting for MICOS-MIB components in control *Mcl1^-/-^* MEFs revealed that while the core MICOS-MIB subunits MIC60 and SAM50 were retained, MIC10 and MIC27, along with MIC13, MIC19 and MTX2 were reduced (Fig. 2E), indicating destabilisation of the assembled MICOS complex following the initiation of apoptosis.

We also sought to identify new IMM proteins that could be important for facilitating IMM herniation and thus focused on the small cluster of mitochondrial proteins with a time dependent increase in enrichment (Fig. 2F & S2C; black box). We identified four IMM proteins that appear to move into proximity with the BAK pore during IMM herniation – 1. the mitochondrial potassium channel subunit *Abcb8* (also known as Mitosur), 2. multifunctional ATPase *Atad3*, 3. phospholipase *Pnlpla8* and 4. the recently uncovered mitochondria localised *Slc35a4* microprotein (SLC35A4-MP) ^(31)^. Given that we observed a temporal increase in enrichment of these four IMM proteins, we hypothesised that they could possess unknown roles in facilitating IMM herniation.

Given that our data demonstrated a loss of MICOS stability during apoptosis, we chose to focus the efforts of further investigation on SLC35A4-MP due to recent suggestions that it could potentially interact with MIC60 ^(32)^. SLC35A4-MP is a highly conserved, upstream open reading frame (uORF)-encoded microprotein, and is the primary translation product of the *Slc35a4* locus ^(32)^. Analysis of SLC35A4 annotated peptides enriched by TurboBAK, enabled us to confirm that the enriched peptides map to the microprotein, and not the canonical SLC35A4 (Fig. S3A). To investigate the function of SLC35A4-MP, we utilised CRISPR/Cas9 to generate a SLC35A4-MP^KO^ in human osteosarcoma (U2OS) cells, and re-expressed SLC35A4-MP^FLAG^ under a doxycycline inducible promoter (Fig. 3A). Western blotting demonstrated that loss of SLC35A4-MP had no effect on levels of BAK, or the MICOS subunits MIC19 and MIC60. Previous studies of SLC35A4-MP had found that the fusion of large peptide tags (such as APEX) disrupted its mitochondrial localisation ^(32)^, thus we initially confirmed that SLC35A4-MP^FLAG^ remained localised to the mitochondria by immunofluorescence assay. Indeed, we observed that SLC35A4-MP^FLAG^ colocalised with the mitochondrial marker TOM20, and formed non-uniform, distinct foci within the mitochondria (Fig. 3B).

**Figure 3:**
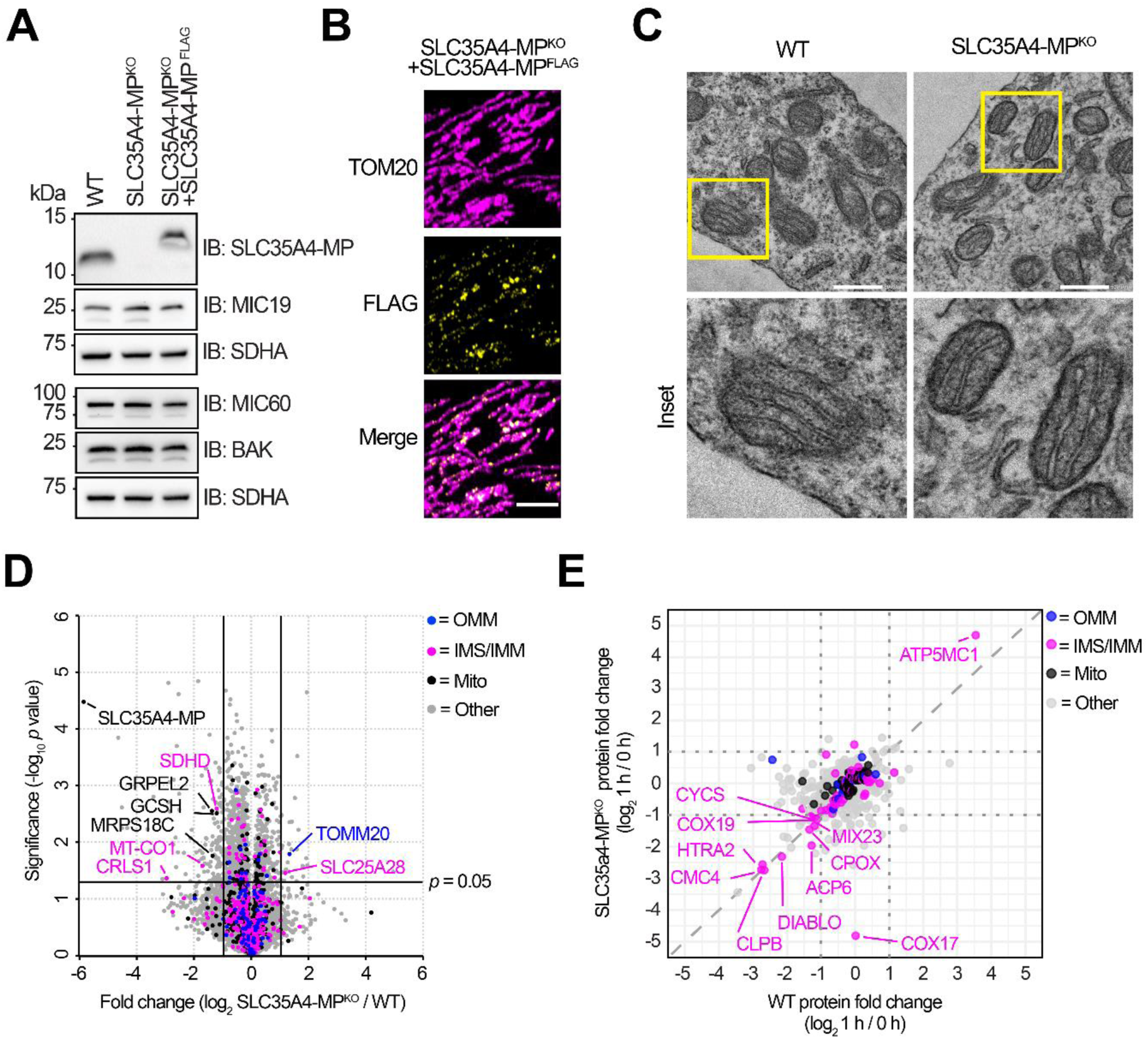
SLC35A4-MP is a mitochondrial microprotein, loss of which does not influence the steady state proteome or cristae architecture. **A**. SDS-PAGE separation and immunoblotting of mitochondria isolated from WT, SLC35A4–MP^KO^ and SLC35A4-MP^KO^ + SLC35A4-MP^FLAG^ U2OS cells. SLC35A4-MP^FLAG^ expression was induced with 5 ng/ml doxycycline for 48 h. SDHA serves as a loading control. Representative of n = 3 biological replicates. **B.** Representative confocal images of SLC35A4-MP^KO^ + SLC35A4-MP^FLAG^ U2OS cells. SLC35A4-MP^FLAG^ expression was induced with 5 ng/ml doxycycline for 48 h. Immunofluorescence was used to label mitochondria (Anti-TOM20, magenta), and determine SLC35A4-MP localisation (anti-FLAG, yellow). Scale bar indicates 5 µm. **C.** Electron microscopy of WT and SLC35A4-MP^KO^ U2OS cells. Scale bars indicate 500 nm. **D.** The steady-state whole cell proteome of SLC35A4-MP^KO^ vs WT. MitoCarta 3.0 proteins are highlighted in black, and those with OMM or IMM localisations are highlighted in blue and pink respectively. **E.** Whole cell proteome of SLC35A4-MP^KO^ vs WT U2OS in apoptotic vs non-apoptotic conditions. The log_2_ (fold change) of significantly altered (*p* < 0.05) WT and KO proteins following 1 h apoptotic treatment was plotted against each other; WT (x-axis) and KO (y-axis). Mitochondrial proteins are highlighted in black, and those with OMM or IMS/IMM localisation are highlighted in blue or pink respectively.

Electron microscopy of SLC35A4-MP^KO^ cells revealed no discernible difference in mitochondrial morphology or cristae architecture at steady state (Fig. 3C). To investigate whether SLC35A4-MP loss had any impact on the whole cell proteome, we performed label free proteomic analysis on whole cell lysate from control and SLC35A4-MP^KO^ cells at steady state (Fig. 3D) and under apoptotic conditions (Fig. 3E). Loss of SLC35A4-MP had minimal impact on the mitochondrial proteome, with only modest changes in a small number of mitochondrial proteins observed. Under apoptotic conditions, both WT and SLC35A4-MP^KO^ cells had a similar reduction in mitochondrial proteins from the intermembrane space (IMS) and IMM, including the pro-apoptotic factors cytochrome *c* (CYCS), HTRA2 and DIABLO (SMAC)^(33–35)^. Additionally, loss of the CIV assembly factor COX17 was observed in SLC35A4-MP^KO^ cells. From this data, we concluded that SLC35A4-MP is not essential for maintaining mitochondrial ultrastructure, nor does the loss of SLC35A4-MP influence changes to the whole-cell proteome at steady-state or during apoptosis.

To further characterise SLC35A4-MP, we performed affinity enrichment mass spectrometry (AE-MS) to identify SLC35A4-MP interacting proteins at steady state (Fig. 4A) and under apoptotic conditions (Fig. 4B). Analysis of the SLC35A4-MP^FLAG^ steady-state interactome revealed strong enrichment of proteins associated with IMM organisation and cristae formation, including known members of the MICOS-MIB and the IMM large GTPase OPA1 (Fig. 4A & S4A). Indeed, the MICOS-MIB was consistently enriched before and after 1 h apoptosis (Fig. S4B), suggesting that this complex is a stable interactor of SLC35A4-MP, and remains associated with SLC35A4-MP during early apoptotic stages. Given the stable interaction between SLC35A4-MP and the MICOS-MIB complex, we investigated the stability of the MICOS complex in mitochondria lacking SLC35A4-MP, however, no differences were observed when compared to control mitochondria (Fig. 4C).

**Figure 4:**
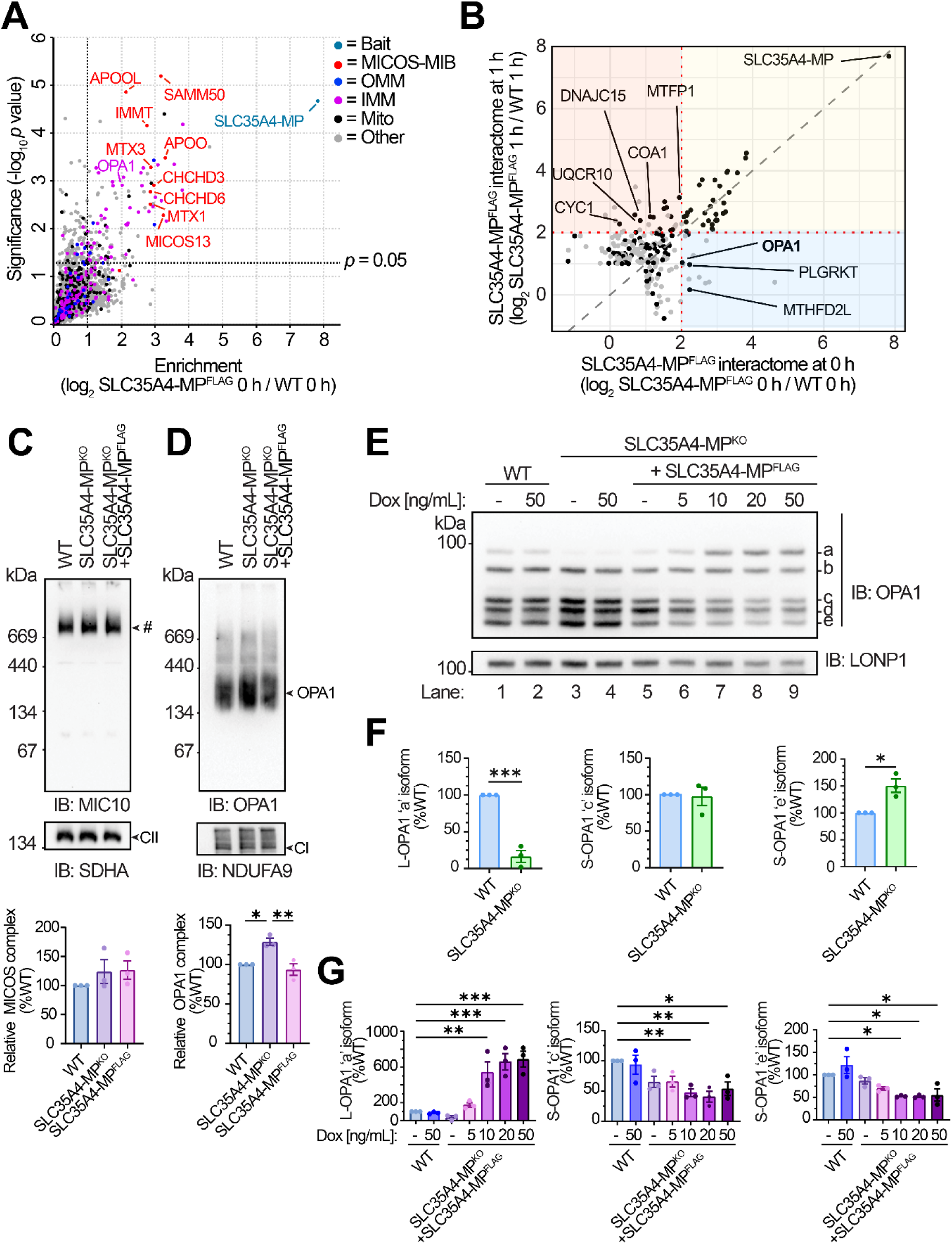
SLC35A4-MP interacts with crucial inner mitochondrial membrane organising proteins, modulates OPA1 oligomerisation and L-OPA1 abundance. **A**. The steady-state SLC35A4-MP interactome. Proteins with a log_2_ (SLC35A4-MP^FLAG^ 0 h / WT 0 h) > 1, *p* value < 0.05, were considered significantly enriched. Proteins with MitoCarta 3.0 annotations have been coloured as indicated **B.** Comparison of log_2_ (SLC35A4-MP^FLAG^ / WT) enrichment of the SLC35A4-MP^FLAG^ interactome at 0 h and 1 h apoptosis. Mitochondrial proteins are coloured in black. Proteins preferentially enriched at 0 h or 1 h post induction of apoptosis are highlighted by the blue and red regions respectively. Stable interactors are highlighted by the yellow region (refer to Fig. S4B) **C-D.** BN-PAGE analysis of 1% digitonin soluble complexes from isolated mitochondria, immunoblotting using antibodies against **C)** MIC10 (MICOS complex) and **D)** OPA1 respectively. NDUFA9 (CI) and SDHA (CII) serve as a loading controls. Representative of n = 3 independent experiments. For densitometric quantification of soluble MIC10 or OPA1 containing complexes, complex abundance was normalised to NDUFA9 or SDHA respectively and displayed relative to WT levels. Data represents n = 3 independent experiments. Error bars indicate mean ± SEM. * = *p* value < 0.05, ** = *p* value < 0.01 by one-way ANOVA with Tukey’s multiple comparison test. **E.** SDS-PAGE analysis of OPA1 isoforms in mitochondria isolated from control, SLC35A4-MP^KO^ and SLC35A4-MP rescue U2OS cells (SLC35A4-MP^FLAG)^. SLC35a4-MP^FLAG^ expression was induced with doxycycline for 48 h (concentrations as indicated). LONP1 serves as a loading control. Representative of n = 3 independent experiments. **F-G.** Densitometric quantification of L-OPA1 ‘a’ isoform, and S-OPA1 isoforms ‘c’ and ‘e’ in **F)** SLC35A4-MP^KO^ (without doxycycline treatment), or **G)** SLC35A4-MP^KO^ cells overexpressing SLC35A4-MP^FLAG^ with increasing concentrations of doxycycline. OPA1 isoform levels were normalised to LONP1 and displayed relative to WT (without doxycycline treatment). n = 3 independent experiments. Error bars indicate mean ± SEM. **F:** *** = *p* value < 0.005 by Student’s t-test. **G:** ** = *p* value < 0.01 *** = *p* value < 0.005 by one-way ANOVA with Tukey’s multiple comparisons test.

At early apoptotic stages (1 h), we observed increased enrichment of 14 mitochondrial proteins including OXPHOS proteins CYC1, COA1 and UQCR10, the IMM chaperone DNAJC15 and the IMM mitochondrial fission process protein 1 MTFP1, a known regulator of IMM dynamics ^(36)^, demonstrating the dynamic nature of the SLC35A4-MP interactome during apoptosis. Only three mitochondrial proteins showed decreased enrichment at 1 h, of which, OPA1 was the only one associated with IMM organisation. This suggested that the interaction between SLC35A4-MP^FLAG^ and OPA1 may be disrupted during apoptosis. To investigate the functional relationship between SLC35A4-MP and OPA1 we explored the effect of SLC35A4-MP loss on soluble OPA1 complexes and OPA1 isoforms. Using BN-PAGE analysis of digitonin-solubilized SLC35A4-MP^KO^ mitochondria, we observed an increase in the abundance of soluble OPA1 complexes in SLC35A4-MP^KO^ mitochondria, that was restored upon complementation with SLC35A4-MP^FLAG^ (Fig. 4D). In WT U2OS cells, five distinct OPA1 isoforms were resolved by SDS-PAGE; “Long” (L-OPA1) isoforms ‘a’ and ‘b’, and “short” (S-OPA1) isoforms ‘c-e’ ^(21)^. Loss of SLC35A4-MP led to a reduction in the L-OPA1 ‘a’ isoform at steady state, and an increase in S-OPA1 ‘e’ isoform (Fig. 4E, compare lanes 1 and 3, Fig. 4F). Meanwhile, total OPA1 levels were unchanged (Fig. S4C) consistent with our whole cell proteomics analysis. Re-expression of SLC35A4-MP^FLAG^ to endogenous levels restored the OPA1 ‘a’ isoform to control levels (Fig. 4E, compare lanes 1 and 6), while over-expression of SLC35A4-MP^FLAG^ resulted in an increased abundance of the ‘a’ isoform, and decreased abundance of S-OPA1 ‘c’ and ‘e’ isoforms in a dose-dependent manner (Fig. 4E, compare lane 1 with lanes 7-9, Fig. 4G). Despite the reduction in L-OPA1 ‘a’ isoform in SLC35A4-MP^KO^ mitochondria at steady state, the ‘a’ isoform remaining was still susceptible to apoptosis-induced cleavage into S-OPA1 isoforms (Fig. S4D-E) and did not influence the overall reduction in OPA1 abundance 1 h post induction of apoptosis. This inverse relationship between SLC35A4-MP abundance and OPA1 ‘a’ isoform processing suggests that SLC35A4-MP loss may specifically influence the proteolytic cleavage of the L-OPA1 ‘a’ isoform.

Given the depleted L-OPA1 ‘a’ isoform in SLC35A4-MP^KO^ cells, we predicted that loss of SLC35A4-MP influences mitochondrial membrane dynamics during apoptosis. To investigate mitochondrial membrane dynamics in real time, we stably expressed TOM20^Halo^, which can be readily visualised by addition of fluorescent Halo-ligands, in our WT, SLC35A4-MP^KO^ and SLC35A4-MP^FLAG^ U2OS cell lines and performed live-cell microscopy. Live-cell imaging revealed a delay in apoptosis-induced mitochondrial fragmentation and mtDNA release in SLC35A4-MP^KO^ cells, that was rescued following re-expression of SLC35A4-MP^FLAG^ (Fig. 5A-C). Notably, delayed mitochondrial fragmentation did not affect cytochrome *c* release in SLC35A4-MP^KO^ cells (Fig. S5A). The delay in both fragmentation and mtDNA release was resolved by 80-100 min post treatment, demonstrated by characteristic TOM20-positive cup-like structures with extruding mtDNA at the 92 min timepoint (Fig. 5A insets), thus indicating that SLC35A4-MP^KO^ mitochondria could still undergo IMM herniation, albeit with delayed kinetics.

**Figure 5:**
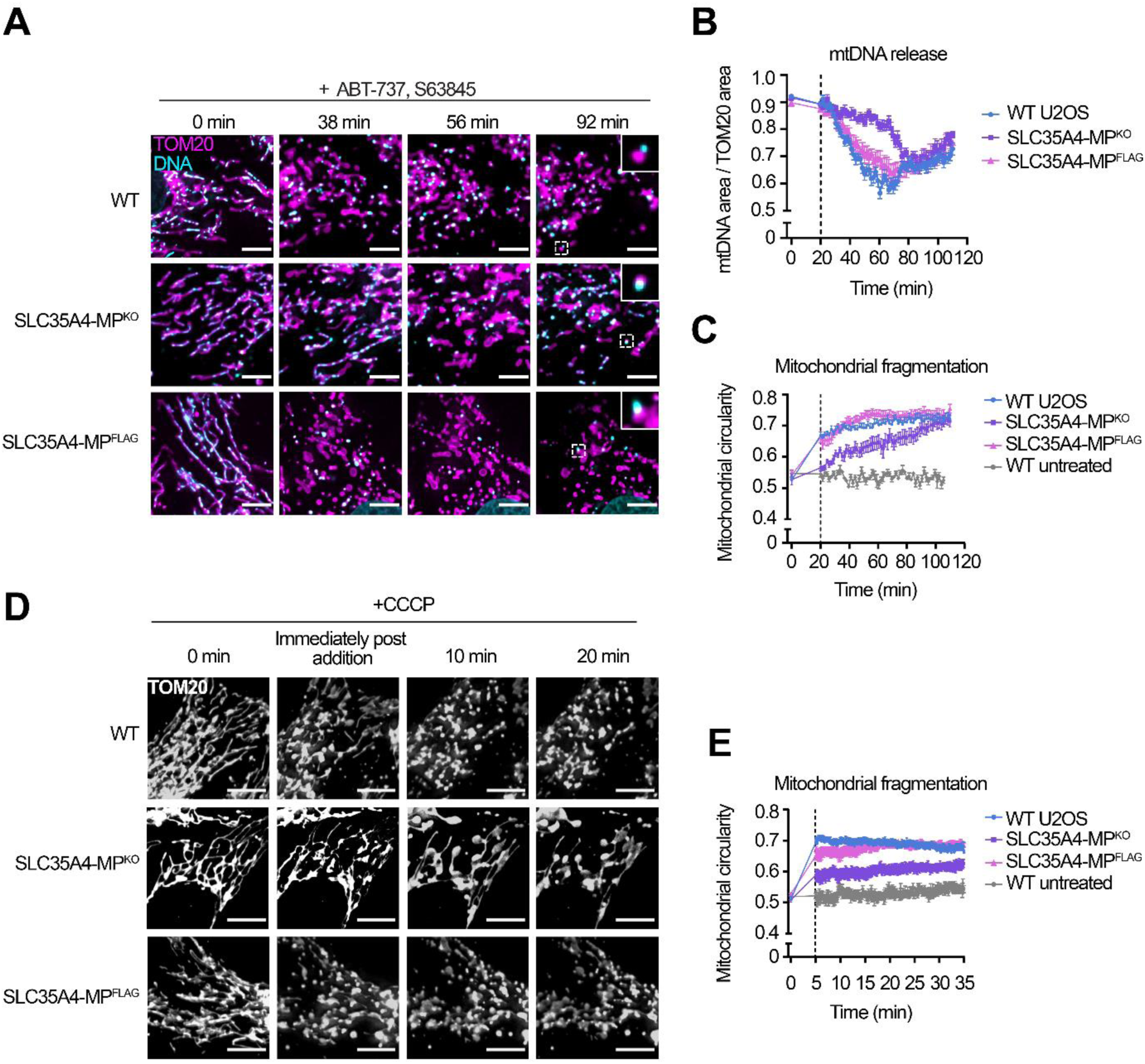
Loss of SLC35A4-MP slows apoptotic-induced mitochondrial fragmentation. **A.** Live-cell imaging of WT, SLC35A4-MP^KO^ and SLC35A4-MP rescue U2OS cells (SLC35A4-MP^FLAG)^ stably expressing TOM20^Halo^ following treatment with ABT-737 [10 µM], S63845 [2 µM] and 1 h pre-treatment with QVD-OPh [20 µM]. Representative images from n=3 independent experiments. Scale bars indicate 5 µm. **B-C.** Quantification of apoptosis induced mtDNA release and mitochondrial fragmentation respectively. Quantification was performed on 10-15 cells from n = 3 independent experiments. Data points indicate mean ± SEM. Dashed line indicates the start of image acquisition **D.** Live cell imaging of SLC35A4-MP^KO^ U2OS cells following CCCP [20µM] treatment. Representative images from n = 3 independent experiments. Scale bars indicate 10 µm. **E.** Quantification of mitochondrial fragmentation. Quantification was performed on 10-15 cells from n=3 independent experiments. Dashed line indicates start of image acquisition.

Given the interaction between SLC35A4-MP and OPA1, we sought to determine whether this delay in mitochondrial network collapse was specific to apoptotic stress, or a broader facet of how SLC35A4-MP^KO^ mitochondria respond to alternate stress conditions. Using either the protonophore, carbonyl cyanide m-chlorophenyl hydrazone (CCCP), a mitochondrial membrane potential uncoupler (Fig. 5D-E) or H_2_O_2_-induced oxidative stress (Fig. S5B-C) we also observed impaired mitochondrial network dynamics SLC35A4-MP^KO^ cells. CCCP treatment induced rapid mitochondrial fragmentation in WT mitochondria however, SLC35A4-MP^KO^ mitochondria resisted CCCP stress and a more reticular network was sustained for 35 mins post treatment. This impairment was specifically dependent on the loss of SLC35A4-MP, since re-expression of SLC35A4-MP^FLAG^ could rescue the phenotype. Taken together, these results indicate that loss of SLC35A4-MP leads to a subtle, but robust delay in mitochondrial fragmentation, not only in response to apoptotic stimulus, but as a general hallmark of the capacity of cells lacking SLC35A4-MP to respond to mitochondrial stress.

## Discussion

Mitochondrial fission and fusion must be delicately balanced and therefore tightly controlled to ensure appropriate mitochondrial network health, function and quality control ^(37, 38)^. Indeed, the control of mitochondrial dynamics is crucial for a multitude of mitochondrial processes including the efficient execution of programmed cell death, which includes network-wide mitochondrial membrane remodelling ^(3, 5, 39–41)^. The intricacies of how integral IMM scaffolding proteins such as the MICOS and OPA1 regulate membrane changes during network-wide mitochondrial remodelling during apoptosis are not fully understood. Here, we have utilised proximity labelling proteomics to capture the dynamic movement of mitochondrial proteins to and from BAK macropores during apoptotic IMM herniation (Fig. 6A). In doing so, we have defined the BAK proximal proteomes at steady state, during apoptosis and IMM herniation. Furthermore, we observed MICOS complex disassembly during apoptosis, which may be required for IMM remodelling events. Indeed, it has been previously reported that a stable MICOS complex is important for preventing mtDNA release and subsequent inflammatory signalling ^(42)^.

**Figure 6.**
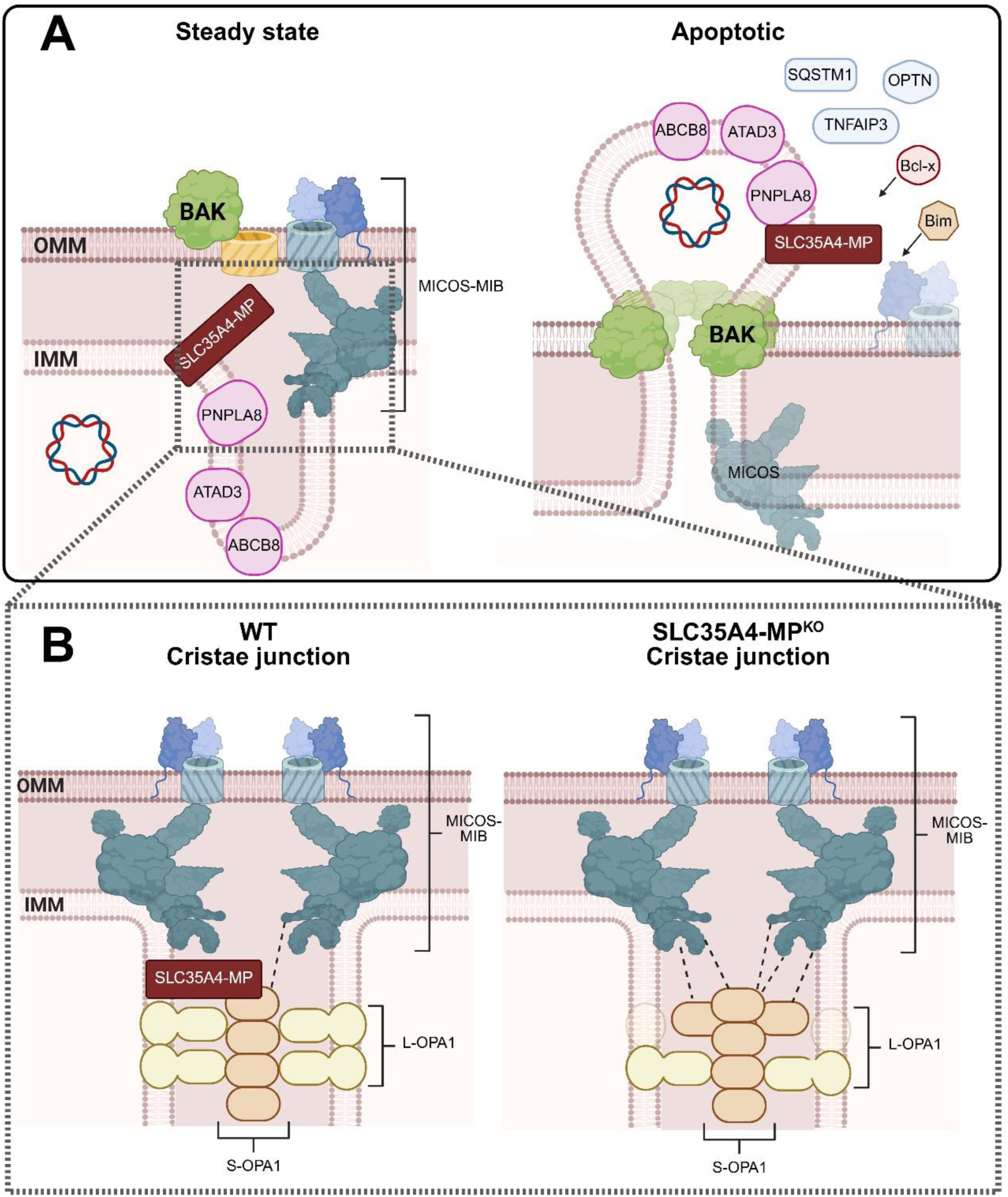
Schematic depicting **A)** the BAK-proximal-proteome under steady-state and apoptotic conditions, and **B)** the influence of SLC35A4-MP loss on cristae junctions.

By extension, this study has also formulated insights into the IMM microprotein SLC35A4-MP. Microproteins originating from uORFs, such as SLC35A4-MP, represent an understudied and potentially underappreciated source of regulators of cellular processes, with a large number of the currently characterized microproteins acting as allosteric modulators of larger proteins ^(43–45)^. Mitochondria in particular are resident to multiple microproteins that regulate crucial processes including the assembly of the electron transport chain ^(46, 47)^ and mitochondrial translation ^(48, 49)^. Our data places SLC35A4-MP at the interface of the MICOS complex and OPA1 at steady-state conditions which, by association, implies a potential role for SLC35A4-MP in regulating mitochondrial dynamics or cristae architecture. Interestingly, we did not observe any changes in cristae or overall network morphology in SLC35A4-MP^KO^ mitochondria at steady-state, indicating that such a role may only be employed under specific conditions of stress. Indeed, our results suggest that SLC35A4-MP dissociates from OPA1 during intrinsic apoptosis and moves into proximity with the BAK pore as IMM herniation progresses.

In line with the association of SLC35A4-MP with OPA1, the loss of SLC35A4-MP induced a reduction in L-OPA1 ‘a’ isoform, and a corresponding increase in the S-OPA1 ‘e’ isoform. Furthermore, we observed increased soluble OPA1 oligomers and delayed mitochondrial fragmentation in response to stresses. L-OPA1 and S-OPA1 isoforms have previously been described to have both redundant and isoform-specific functions ^(20)^. For example, global loss of OPA1 leads to fragmentation of the mitochondrial network, disorganisation of cristae structure, reduced mitochondrial membrane potential and reduced respiratory capacity ^(50–52)^. Meanwhile, L-OPA1, but not S-OPA1, is capable of performing mitochondrial fusion while re-expression of any single OPA1 isoform can completely recover the functional energetic deficiencies associated with OPA1 loss, independently of mitochondrial network morphology ^(53)^. Furthermore, soluble S-OPA1, but not L-OPA1 has been shown to directly bind MIC19 and MIC60 and play a role in maintaining stable cristae junctions ^(54)^. Since we did not observe any changes in cristae architecture related to loss of SLC35A4-MP, we postulate that the accessory role of SLC35A4-MP in modulating OPA1 function is independent of its role in maintaining IMM architecture at steady state. Overall, our data suggests that SLC35A4-MP influences the steady-state abundance of the L-OPA1 ‘a’ isoform and plays a role in modulating mitochondrial fragmentation induced by a variety of stressors. Furthermore, we posit that the resulting increase in soluble S-OPA1 isoforms may result in more rigid cristae junctions that are more resistant to changes in morphology (Fig. 6B).

How SLC35A4-MP loss alters the steady-state abundance of OPA1 isoforms is not clear. In humans and mice, the IMS-residing metalloprotease OMA1 and i-AAA protease YME1L cleave L-OPA1 to constitutively maintain an appropriate balance of L-OPA1 and S-OPA1 isoforms at steady-state. OMA1-mediated degradation of L-OPA1 occurs in response to a variety of insults that lead to fragmentation of the mitochondrial network ^(21, 22)^. In addition, the rhomboid protease PARL is involved in the production of soluble OPA1. Furthermore, PARL expression can modulate cytochrome *c* release and OPA1-dependent cristae remodelling during apoptosis ^(55)^. Understanding whether SLC35A4-MP protects the L-OPA1 ‘a’ isoform from proteolytic processing by any of these proteases could provide important mechanistic insight into the regulation of OPA1 isoform abundance, since loss of SLC35A4-MP specifically causes depletion of the L-OPA1 ‘a’isoform. Intriguingly, both PARL and YME1L, but not OMA1, were significantly enriched by AE-MS in SLC35A4-MP^FLAG^-expressing cells following apoptotic treatment, however, the functional relationship between these proteases and SLC35A4-MP remains to be elucidated.

Whilst our study investigated the role of SLC35A4-MP in mitochondrial dynamics during mitochondrial stress, including apoptosis, recent findings show that loss of SLC35A4-MP *in vivo* leads to altered lipid metabolism and increased mitochondrial size in murine brown adipose tissue (BAT) ^(32)^. Furthermore, *Slc35a4-mp^-/-^* mice were more sensitive to acute cold exposure compared to control mice. Intriguingly, BAT specific *Opa1^-/-^* mice have also been shown to have reduced lipid synthesis and lipolysis signalling, along with an impaired response to cold exposure ^(56)^. The influence of SLC35A4-MP on OPA1, and the perturbation of mitochondrial dynamics during stress in the absence of SLC35A4-MP, may provide a mechanistic link between the *Slc35a4-mp^-/-^*and *Opa1^-/-^* phenotypes – although the BAT-specific influence of SLC35A4-MP on OPA1 isoforms is yet to be investigated.

Our characterisation of SLC35A4-MP has revealed a potentially important function in modulating OPA1 and mitochondrial dynamics, however, it does not appear to play a primary role in controlling apoptotic IMM herniation. Fission/fusion incompetent MEF cells (*Drp1^-/-^*, *Mfn1/2^-/-^*, *Opa1^-/-^*) are still capable of releasing cytochrome *c,* and subsequently mtDNA, following apoptotic stimuli ^(3)^. Intriguingly, in addition to SLC35A4-MP, PNPLA8 and ATAD3 were also identified as IMM proteins that come into proximity with the BAK pore during herniation. PNPLA8 is a phospholipase responsible for the release of oxidised cardiolipin, a key lipid component of the IMM, from membranes ^(57)^ and could conceivably play a role in mediating IMM permeability during herniation. In contrast, ATAD3 has previously been shown to bind mtDNA nucleoids and is required for upholding cristae structure ^(58–60)^. Furthermore, ATAD3 directly interacts with SAM50, and the ATAD3-SAMM50 axis is necessary for the extraction of mitochondrial nucleoids for delivery to endosomes following mtDNA damage ^(61)^. The established relationship between ATAD3, mtDNA organisation and cristae structure, and its newly identified proximity to the BAK pore during IMM herniation places ATAD3 as a prime candidate for further investigation into proteins required for IMM remodelling during apoptosis.

In conclusion, our study describes the BAK-proximal-proteome at steady state and during apoptosis. We have demonstrated a dynamic interaction between the recently identified IMM microprotein SLC35A4-MP and OPA1 and demonstrated an inverse relationship between SLC35A4-MP expression and the abundance of long OPA1 ‘a’ isoform. We posit that SLC35A4-MP may play a previously unappreciated role in mitochondrial membrane dynamics, affecting the timely responsiveness of mitochondrial networks during stress conditions.

## Methods

### Cell culture and genome editing

Immortalised mouse embryonic fibroblasts (MEFs) were previously generated: *Mcl1*^-/-^ (a gift from B.Kile) and *Mcl1*^-/-^*Bak*^-/-^*Bax*^-/-^ (a gift from D. Huang). Human osteosarcoma cells (U2OS) were purchased from the ATCC, and a clonal line was established. MEF cells were cultured in DMEM high glucose media, supplemented with 10% (v/v) foetal bovine serum (FBS), 1 x penicillin / streptomycin (Sigma-Aldrich; P4333), 1 x Glutamax (Thermo Fisher Scientific; 35050061) and 50 µg / mL uridine (Sigma-Aldrich; U3750). U2OS cells were cultured in RPMI 1640 media, supplemented with the same additions. All cells were grown at 37°C with a humidified 5% CO2 atmosphere.

Gene editing of *Slc35a4-mp* was performed using the pSp-Cas9(BB)-2A-GFP (PX458) CRISPR/Cas9 construct (a gift from F. Zhang; Addgene #48138) ^(62)^, as previously described ^(63)^. A two CRISPR targeting approach to gene disruption was employed, with single guide RNAs (sgRNAs) targeting the 5’ UTR (target sequence: 5-CGAGTGAGATCATCGCGCCCCGG-3), and exon 3 of *Slc35a4-mp* (target sequence: 5-CCGACTAGCTCTAGGTGCCATGG-3) designed using CHOPCHOP ^(64)^. Cells were transfected using Lipofectimine LTX (Thermo Fisher Scientific; 15338100) according to manufacturer’s instructions and clonal populations were isolated by fluorescence-activated cell sorting (FACS; Flowcore, Monash University). SLC35A4-MP loss was confirmed by SDS-PAGE and Western blotting.

For the stable expression of HA-TurboID-BAK, cDNAs for TurboID and BAK were amplified from template plasmids by PCR and cloned into the retroviral vector pBMN-Z (Addgene #1734, re-engineered by removal of the LacZ locus and incorporation of an IRES and puromycin resistance cassette). The N-terminal HA tag was incorporated using an in-frame HA epitope at the 5’end (after the start codon) of the TurboID forward primer. For inducible expression of SLC35A4-MP^FLAG^, SLC35A4-MP cDNA was amplified by PCR from a HEK293T cDNA library first generated using the Superscript III First strand cDNA synthesis kit (Thermo Fisher Scientific; 18080051) using a reverse primer incorporating the sequence for an in-frame FLAG epitope at the 3’ end. SLC35A4-MP^FLAG^ cDNA was cloned into the doxycycline inducible lentiviral vector pLVX-TetOne-Puro (Clontech; 631847). All cloning was performed using Gibson Assembly (NEB), and newly generated plasmids were sequence verified (Micromon, Monash University). For the stable expression of TOMM20^Halo^, the pMIH-TOM20-HaloTag hygromycin-selectable retroviral vector ^(3)^ was stably expressed in U2OS cells. Viral supernatants were prepared by co-transfecting HEK293T cells with either pBMN-Z, pLVX-TetOne-Puro or pMIH plasmids encoding the cDNA of interest with helper vectors encoding *gag*, *pol*, and *env* genes necessary for viral particle formation and replication. Viral supernatants were collected after 48 h and filtered with a 0.45 µm low protein binding PVDF filter (Merck; SLHV033RS). Target cells were then infected with the relevant viral supernatant, along with 8 µg / mL polybrene (Sigma-Aldrich; H9268). Transduced cells were selected with the appropriate selection reagent (pBMN-Z, pLVX-TetOne-Puro: 2 µg / mL puromycin, pMIH: 250 μg / mL Hygromycin B Gold (Invivogen NC9163750). For HA-TurboID-BAK and SLC35A4-MP^FLAG^ expressing cells, a clonal population was isolated by FACS (Flowcore, Monash University).

### Induction of apoptosis

Apoptosis was induced in *Mcl1*^-/-^ MEFs by treatment with 2µM ABT-737 for the indicated duration. Unless otherwise stated, caspases were inhibited by pre-treatment with 20µM QVD-OPh (MedKoo Biosciences; #526845) for 1 h prior to treatment with ABT-737. For human U2OS cells, apoptosis was induced with 10µM ABT-737 and 2µM of the *Mcl1*^-/-^ inhibitor S63845 (Cayman Chemical; #21131).

### Mitochondrial isolation and sub-cellular fractionation

Mitochondria were isolated by differential centrifugation as previously described ^(65, 66)^. The protein content of isolated mitochondria, or whole cell suspensions were quantified by bicinchoninic acid assay (BCA; Thermo Fisher Scientific; 23227). For subcellular fractionation, harvested cells were permeabilised with 0.0125% digitonin prepared in lysis buffer (20 mM HEPES KOH pH 7.5, 93 mM sucrose, 100mM KCl, 2.5 mM MgCl_2_,) at 4°C for 10 min. A half-volume ‘Total’ fraction was taken and quantified by BCA. The pellet and supernatant fractions were separated by centrifugation at 13,000 *g*, 4°C for 10 min.

### Polyacrylamide gel electrophoresis and Western blotting

SDS-PAGE using a tris-tricine buffered system and BN-PAGE were performed as previously described ^(67–69)^. For SDS-PAGE, samples were solubilised in 1 x Laemmli buffer supplemented with 100 mM DTT and heated at 95°C for 5 min. Samples were separated with either 8%, 10% or continuous 10-16% acrylamide gradient gels depending on the proteins of interest. For BN-PAGE, samples were solubilised in a detergent buffer (20 mM Bis-Tris pH 7.0, 50 mM NaCl, 10% (w/v) glycerol) containing 1 % digitonin for 10 min, followed by centrifugation at 16,000 *g,* 4 °C for 10 min. 10x BN-PAGE loading dye (5% (w/v) Coomassie Brilliant Blue G250, 500 mM ε-amino n-caproic acid, 100 mM Bis-Tris pH 7.0) was added at a 1:10 dilution to the clarified supernatant, and loaded onto a 4-13% continuous acrylamide gradient gel. Following electrophoresis, proteins were transferred onto a PVDF membrane (Merck; IPVH00010), or for sub-cellular fractionation, Immobilon-PSQ PVDF (Merck; ISEQ00010) using either a Novex Semi-Dry Blotter (Thermo Fisher Scientific) or an Invitrogen Power Blotter System (Thermo Fisher Scientific) according to manufacturer’s instructions.

Primary antibodies used in this study are listed in Supplementary Table 1. Anti-mouse (Sigma-Aldrich; A9044) or anti-rabbit (Sigma-Aldrich; A0545) horseradish peroxidase-conjugated secondary antibodies were used at a dilution of 1:10,000. Clarity Western ECL chemiluminescent substrate (BioRad; 1705061) was used for detection on a BioRad ChemiDoc XRS+ imaging system according to manufacturer’s instructions.

Densitometry was performed in Image Lab™ Software (BioRad; version 6.0.1) by taking a defined sized region around each band/complex of interest and measuring the pixel intensity relative to WT at baseline. To account for background, the pixel intensity of an additional region of the same size was selected outside of the regions of interest and subtracted from the regions of interest. Protein bands/complexes were normalised relative to the pixel intensity of the associated loading control and shown as a percentage relative to control. Data analysis and statistical tests were performed in GraphPad Prism (version 10.1.2).

### Caspase-Glo assay

Caspase activity was determined using the Caspase-Glo 3/7 luminescence assay (Promega; #G8093) and was performed as per the manufacturer’s instructions, with modifications. In brief, 5000 cells were seeded in triplicate into a 96 well plate 24 h prior to performing the assay. Media was replaced 1 h prior to treatment with 2µM ABT-737 for the time periods indicated. QVD-OPh was not added. Between treatments, plates were returned to 37°C with a humidified 5% CO_2_ atmosphere. Following treatment, an equal volume of Caspase-Glo reagent was added, mixed, and incubated in the dark for 30 min. Samples were transferred to a white opaque 96 well plate and luminescence was read on either FLUOstar Optima or CLARIOstar plate reader (BMG Labtech). Luminescence was graphed relative to the no treatment (0 h) control for each experiment using GraphPad Prism (version 10.1.2). Statistical significance was determined by two-way ANOVA with Tukey’s multiple comparison test.

### Immunofluorescence Assay and Fixed-Cell Imaging

Protein detection by immunofluorescence was performed as previously described with some modifications ^(70)^. MEFs were seeded onto Histogrip-coated #1.5 glass cover slips (Zeiss Microscopy). To induce herniation, MEF cells were pre-treated with [20 µM] QVD-OPh for 1 h prior to treatment with ABT-737 [2 µM]. Cells were fixed with pre-warmed (37°C) 4% (w/v) paraformaldehyde in 1 × PBS for 15 min before washing three times with 1 × PBS. Samples were then permeabilised with 0.1% (v/v) Triton X-100 in 1 × PBS for 10 minutes at room temperature before being washed three times with 1 × PBS followed by blocking in 3 % (w/v) BSA in 1 × PBS for 30 min. Samples were incubated with desired primary antibodies for 2 h at room temperature (Supplementary table 1). Samples were then washed three times with 1X PBS and incubated with anti-rabbit-IgG Alexa Fluor™-647 (ThermoFisher Scientific; #A-21244), anti-mouse-IgM Alexa Fluor™-488 (ThermoFisher Scientific; #A-21042), and anti-mouse-IgG1 Alexa Fluor™-555 (ThermoFisher Scientific; #A-21127) secondary antibodies for 1 h. Samples were washed a final three times before coverslips were mounted onto microscope slides with DABCO mounting medium (1,4-Diazobicyclo-(2,2,2-octane) (DABCO) in 90% (v/v) glycerol, Tris-Cl, pH 8.0) and sealed with nail polish.

Samples were imaged on the Zeiss LSM 980 confocal microscope equipped with 405, 445, 488, 514, 561, and 639 nm lasers and Airyscan 2 detector, 32+2 spectral GaAsP detector with two flanking PMTs and transmitted light PMT TLD. Confocal images were acquired using a 63x/1.4 NA oil immersion objective in confocal mode using ZEN software (Zeiss; version 3.9). Images were acquired as z-stacks (10 slices) with 0.23 µm step size. The same settings were applied for all experimental groups. All images were processed and converted to maximum intensity projections using FIJI/ImageJ ^(71)^.

### Live cell imaging

U2OS cells were grown in 8-well chambered #1.5 polymer ibiTreat slides (Ibidi; #80826), and incubated with 50 nM Janelia Fluor JFX650 HaloTag-specific dye ^(72)^ or 1 × SYBR Gold Nucleic Acid Stain (ThermoFisher Scientific; #S11494) for 30 min at 37 °C and 5 % CO_2_. For imaging of apoptotic cells, cells were then incubated with 20 μM QVD-OPh at 37°C and 5% CO_2_ for 1 h. For all live-cell imaging experiments, media was then replaced with Leibovitz’s L-15 medium (ThermoFisher Scientific; #21083027) supplemented with 10 % (v/v) FBS, 25 mM HEPES (ThermoFisher Scientific; #15630080), 10 μl / mL penicllin / streptomycin (Sigma-Aldrich; #P4333), and 1 × GlutaMAX (ThermoFisher Scientific; #35050061). For non-oxidative stress experiments, 20 μM TROLOX (Santa Cruz Biotechnology; #53188-07-1) was also included in the media. Cells were imaged on an inverted Leica DMi8 Thunder Deconvolution widefield microscope in a 37 °C environment chamber using a 100x /1.4 oil objective. Fluorescence emission was detected by a 4.2 MP sCMOS K8 camera, with exposure kept below 200 ms. Z-stacks were acquired with a 0.27 μm step size (8-10 slices), and images shown are maximum-intensity projections. Following imaging of cells at baseline, cells were treated with either apoptotic-inducing drugs ABT-737 [10 μM] and S63845 [2 μM], CCCP [20 μM] (Sigma-Aldrich; #C2759), or H_2_O_2_ [500 μM] (Sigma-Aldrich; #1072090500) before capturing time-lapse images at 90 sec intervals for 90 min, 5 sec intervals for 30 min or 90 sec intervals for 2 h respectively, with Leica Microsystems SVCC Deconvolution algorithm applied. All image data was analysed using the FIJI distribution of ImageJ ^(71)^. Mitochondrial morphology was quantified using FIJI’s in-built circularity measures, and mtDNA release was measured by using FIJI’s JACoP colocalisation plugin to quantify the percentage overlap between masks of the mitochondria and mtDNA channels ^(71)^.

### Transmission Electron Microscopy

Electron microscopy was performed on U2OS cells by the Monash Ramaciotti Centre for Cryo-Electron Microscopy as previously described ^(73)^. All steps were carried out using the PELCO BioWave system (Ted Pella, Inc.). Ultrathin sections (70 nm) were cut using a Leica UC7 ultramicrotome (Leica Microsystems) and imaged at 80 kV on a JEOL1400 Plus TEM equipped with a high sensitivity bottom mount CMOS ‘Flash’ camera.

### Mass spectrometry: Proximity labelling

To perform biotin labelling, TurboID-BAK expressing MEFs were treated with 2 µM ABT-737 and 20 µM QVD-OPh (1 h pre-treat) for the desired time points, in triplicate. Subsequently, 50 µM biotin was added for 10 min. Cells were harvested with ice -cold 1 x PBS and quantified by BCA assay. 500 µg of whole cell material was solubilised in RIPA lysis buffer (50 mM Tris-HCl pH 7.5, 150 mM NaCl, 1 mM EDTA, 1 mM EGTA, 1% NP-40, 0.1% SDS) supplemented with 1 x protease inhibitor cocktail (Sigma-Aldrich; #11873580001), 0.5% sodium deoxycholate and 0.125 U/ml Turbonuclease (Signa-Aldrich; #T4330). Biotinylated proteins were enriched with streptavidin-coated magnetic beads (Thermo Fisher Scientific; #88816), with subsequent washes with TAP lysis buffer (50 mM HEPES-KOH pH 7.6, 100 mM KCl, 10 % glycerol, 2 mM NP-40) and 50 mM ammonium bicarbonate (pH 8.0) to remove unbound proteins. Captured proteins were reduced and alkylated with 10 mM TCEP (Thermo Fisher Scientific; #77720) and 40 mM iodoacetamide respectively, before being subjected to overnight digest with 2.5 µg / ml mass spectrometry grade trypsin (Promega; V5280). Following digest, desalting was performed using in-house-generated Styrene Divinyl Benzene (SDB-XC) (Empore; #66884-U) StageTips ^(74)^. Samples were dried and resuspended in 12 µL 2% ACN, 0.1% FA (v/v) containing iRT peptides (Biognosys, 1:100 ratio) for LC-MS/MS analysis.

### Whole cell proteomics

Whole cell proteomics was performed on 150 µg of U2OS WT or SLC35A4-MP^KO^ whole cell material, at either 0 h or 1 h post apoptotic induction as previously described. Samples were prepared in triplicate. Sample preparation was performed with S-Trap mini columns (Protifi: CO2-mini-80) as per the manufacturer’s instructions. Proteins were digested with mass spectrometry grade trypsin gold (Promega: V5280) for 16 h, at 37 °C, at a ratio of 1:50 trypsin:protein. Following digest, desalting was performed using in-house-generated Styrene Divinyl Benzene (SDB-XC) StageTips. Samples were dried and resuspended in 15 µL 2% (v/v) ACN, 0.1% FA (v/v) containing iRT peptides (Biognosys, 1:100 ratio) for LC-MS/MS analysis.

### SLC35A4-MP affinity enrichment mass-spectrometry (AE-MS)

Affinity enrichment mass-spectrometry (AE-MS) was performed using 500 µg of U2OS WT or SLC35A4-MP^FLAG^ whole cell material, at either 0 h or 1 h post apoptotic induction as previously described. SLC35A4-MP^FLAG^ expression was induced with 5 ng/ml doxycycline for 48 h. Samples were prepared in triplicate and were solubilised in lysis buffer (20 mM Tris pH 7.4, 50 mM NaCl, 10% glycerol, 100 mM EDTA) containing 1 % (v/v) digitonin for 30 min before centrifugation at 16,000 *g* for 10 min. The clarified supernatant was incubated with ANTI-FLAG M2 affinity gel (Sigma Aldrich; A2220) for 2 h, before washed 30 x with lysis buffer containing 0.1 % (v/v) digitonin via a vacuum manifold system. Following washing, bound proteins were eluted with 150 µg / ml DYKDDDDK tag peptide (Assay Matrix; A6002). Eluted proteins were precipitated with five volumes of ice-cold acetone, prior to overnight storage at -20°C. Acetone was removed, and samples were prepared for mass spectrometry with S-Trap mini columns (Protifi: CO2-mini-80), and StageTip desalting. Proteins were digested for 16 h, at 37°C with 4 µg trypsin gold per sample. Dried peptides were resuspended in 12 µL 2 % ACN, 0.1 % FA (v/v) containing iRT peptides (Biognosys, 1:100 ratio) for LC-MS/MS analysis.

### Instrumentation

For LC-MS/MS analysis, 1 µl of the resuspended sample was loaded onto either an Orbitrap Exploris 480, or Orbitrap Eclipse Tribrid Mass Spectrometer coupled to a Vanquish Neo UHPLC (Thermo Fisher Scientific). Samples were loaded at a flow rate of 15 µL/min onto an Acclaim PepMap 100 trap column (100 μm x 2 cm, nanoViper, C18, 5 μm, 100Å; Thermo Fisher Scientific) which was maintained at 40 °C. Peptides were eluted from the trap column at a flow rate of 0.25 µL/min through the Acclaim PepMap RSLC analytical column (75 μm x 50 cm, nanoViper, C18, 2 μm, 100Å; Thermo Fisher Scientific). The HPLC gradient was set to 158 min. Buffer A was 0.1% FA, buffer B was 80% ACN (v/v), 0.1% FA. The gradient started at 2.5% B before reaching 7.5% B after 3 min, 37.5% B after 123 min, 42.5% B after 126 min and 99% B after 131 min before dropping to 2.5% B at 138 min, for the remainder of the run. The mass spectrometer was operated in a data-independent mode (DIA). MS/MS was run with FAIMS. FAIMS compensation voltages (CVs) included -45 V and -60 V. The MS1 Orbitrap resolution for -45 V and -60 V were set to 60K and 120K, respectively. A full ms1 scan (normalised AGC target: 300%; scan range: 380-1000 m/z) followed by sequential DIA windows (isolation width: 24 m/z) were acquired (scan range: 145 – 1450 m/z; resolution: 30 000; normalised AGC target: 1000%).

### Data processing

DIA analysis was performed using Spectronaut (Biognosys, Version 20.2) using a Direct DIA analysis approach. Spectra were searched against either the *Mus Musculus* (UP000000589) or *Homo sapiens* (UP000005640) UniProt FASTA database. Enzyme specificity was set at Trypsin/P, the digest type was specific, with 2 missed cleavages allowed, a minimum peptide length of 7 and a maximum peptide length set at 52. The imputing strategy was set as global. Oxidation of methionine and protein N-terminal acetylation were set as variable modifications. Carbamidomethylation of cysteines was set as a fixed modification. All other settings were left as default. Following identification and quantification of peptides, raw intensities were exported to Perseus (version 1.6.15) for filtering of known contaminants, GO term and MitoCarta 3.0 annotation and statistical analysis. *P*-values were determined using a two-sided Students’ t-test. For proximity labelling and AE-MS, proteins with a log_2_ (fold change) > 1 and *p* value < 0.05 were considered significantly enriched. For whole cell proteomics, proteins with a log_2_ (fold change) > 1 or < 1 and *p* value < 0.05 were considered significantly perturbed. Heatmaps were generated using Morpheus (https://software.broadinstitute.org/morpheus). Hierarchical clustering was performed using one-minus Pearson correlation. Protein-protein network analysis and functional GO term enrichments were performed with StringDB. ^(75)^. Protein-protein connections indicate high confidence interactions (confidence score > 0.70, sourced from textmining, experimental evidence, databases, co-expression and co-occurrence). Protein networks were clustered using Cytoscape v3.10.2, using the ClusterOne algorithm.

## Acknowledgements

We acknowledge funding from the Australian Research Council (DP190103068 (M.T.R), DP220103559 (M.T.R. & K.M.)) and National Health and Medical Research Council (Peter Doherty ECF GNT1161352 (K.M.), EL1 Investigator Grant GNT2010149 (L.E.F.)). We thank the Monash Proteomics and Metabolomic Platform (MPMP) for provision of instrumentation, training and technical assistance for all mass spectrometry experiments, the Monash Micro-Imaging facility (MMI) for the provision of instrumentation, training and technical assistance for all fixed and live cell imaging experiments and we thank the Monash Ramaciotti Centre for Cryo Electron Microscopy for their provision of instrumentation and technical assistance for all transmission electron microscopy experiments. In addition, we thank the Monash Flowcore and Micromon platforms for cell sorting and Sanger sequencing services.

## Author contributions

**Matthew P. Challis:** Conceptualization, data curation, formal analysis, investigation, methodology, project administration, visualization, writing – original draft

**Stephanie M. Molé:** Formal analysis, investigation, methodology, visualisation

**Saveen Giri:** Formal analysis, investigation, methodology, visualisation

**Ryker Dumbrill:** Formal analysis, investigation, methodology

**Matthew J. Eramo:** Conceptualization, investigation, methodology

**Alice J. Sharpe:** Investigation, methodology

**Sarah E. J. Morf:** Investigation, methodology, writing – reviewing and editing

**Kate McArthur:** Funding acquisition, investigation, methodology, project administration, resources, supervision, writing-review & editing

**Luke E. Formosa:** Funding acquisition, investigation, methodology, project administration, resources, supervision, visualisation, writing-review & editing

**Michael. T. Ryan:** Conceptualization, funding acquisition, project administration, resources, supervision, visualization, writing-review & editing

## References

1. Youle RJ, Strasser A. The BCL-2 protein family: opposing activities that mediate cell death. Nat Rev Mol Cell Biol. 2008;9(1):47–59. 10.1038/nrm2308

2. Glover HL, Schreiner A, Dewson G, Tait SWG. Mitochondria and cell death. Nat Cell Biol. 2024. 10.1038/s41556-024-01429-4

3. McArthur K, Whitehead LW, Heddleston JM, Li L, Padman BS, Oorschot V, et al. BAK/BAX macropores facilitate mitochondrial herniation and mtDNA efflux during apoptosis. Science. 2018;359(6378). 10.1126/science.aao6047

4. Frank S, Gaume B, Bergmann-Leitner ES, Leitner WW, Robert EG, Catez F, et al. The Role of Dynamin-Related Protein 1, a Mediator of Mitochondrial Fission, in Apoptosis. Dev Cell. 2001;1(4):515–25. 10.1016/S1534-5807(01)00055-7

5. Riley JS, Quarato G, Cloix C, Lopez J, O’Prey J, Pearson M, et al. Mitochondrial inner membrane permeabilisation enables mtDNA release during apoptosis. EMBO J. 2018;37(17). 10.15252/embj.201899238

6. White MJ, McArthur K, Metcalf D, Lane RM, Cambier JC, Herold MJ, et al. Apoptotic caspases suppress mtDNA-induced STING-mediated type I IFN production. Cell. 2014;159(7):1549–62. 10.1016/j.cell.2014.11.036

7. Ichim G, Lopez J, Ahmed SU, Muthalagu N, Giampazolias E, Delgado ME, et al. Limited mitochondrial permeabilization causes DNA damage and genomic instability in the absence of cell death. Mol Cell. 2015;57(5):860–72. 10.1016/j.molcel.2015.01.018

8. Victorelli S, Salmonowicz H, Chapman J, Martini H, Vizioli MG, Riley JS, et al. Apoptotic stress causes mtDNA release during senescence and drives the SASP. Nature. 2023;622(7983):627–36. 10.1038/s41586-023-06621-4

9. Rongvaux A, Jackson R, Harman CC, Li T, West AP, de Zoete MR, et al. Apoptotic caspases prevent the induction of type I interferons by mitochondrial DNA. Cell. 2014;159(7):1563–77. 10.1016/j.cell.2014.11.037

10. Gkirtzimanaki K, Kabrani E, Nikoleri D, Polyzos A, Blanas A, Sidiropoulos P, et al. IFNα Impairs Autophagic Degradation of mtDNA Promoting Autoreactivity of SLE Monocytes in a STING-Dependent Fashion. Cell Rep. 2018;25(4):921–33.e5. 10.1016/j.celrep.2018.09.001

11. Matsui H, Ito J, Matsui N, Uechi T, Onodera O, Kakita A. Cytosolic dsDNA of mitochondrial origin induces cytotoxicity and neurodegeneration in cellular and zebrafish models of Parkinson’s disease. Nat Commun. 2021;12(1):3101. 10.1038/s41467-021-23452-x

12. Hancock-Cerutti W, Wu Z, Xu P, Yadavalli N, Leonzino M, Tharkeshwar AK, et al. ER-lysosome lipid transfer protein VPS13C/PARK23 prevents aberrant mtDNA-dependent STING signaling. J Cell Biol. 2022;221(7). 10.1083/jcb.202106046

13. Strauss M, Hofhaus G, Schröder RR, Kühlbrandt W. Dimer ribbons of ATP synthase shape the inner mitochondrial membrane. Embo J. 2008;27(7):1154–60. 10.1038/emboj.2008.35

14. Barbot M, Jans Daniel C, Schulz C, Denkert N, Kroppen B, Hoppert M, et al. Mic10 Oligomerizes to Bend Mitochondrial Inner Membranes at Cristae Junctions. Cell Metab. 2015;21(5):756–63. 10.1016/j.cmet.2015.04.006

15. Daumke O, van der Laan M. Molecular machineries shaping the mitochondrial inner membrane. Nat Rev Mol Cell Biol. 2025;26(9):706–24. 10.1038/s41580-025-00854-z

16. Golla VK, Boyd KJ, May ER. Curvature sensing lipid dynamics in a mitochondrial inner membrane model. Commun Biol. 2024;7(1):29. 10.1038/s42003-023-05657-6

17. Eramo MJ, Lisnyak V, Formosa LE, Ryan MT. The ‘mitochondrial contact site and cristae organising system’ (MICOS) in health and human disease. J Biochem. 2019;167(3):243–55. 10.1093/jb/mvz111

18. Stephan T, Brüser C, Deckers M, Steyer AM, Balzarotti F, Barbot M, et al. MICOS assembly controls mitochondrial inner membrane remodeling and crista junction redistribution to mediate cristae formation. EMBO J. 2020;39(14):e104105. 10.15252/embj.2019104105

19. Frezza C, Cipolat S, Martins de Brito O, Micaroni M, Beznoussenko GV, Rudka T, et al. OPA1 Controls Apoptotic Cristae Remodeling Independently from Mitochondrial Fusion. Cell. 2006;126(1):177–89. 10.1016/j.cell.2006.06.025

20. Olichon A, ElAchouri G, Baricault L, Delettre C, Belenguer P, Lenaers G. OPA1 alternate splicing uncouples an evolutionary conserved function in mitochondrial fusion from a vertebrate restricted function in apoptosis. Cell Death Differ. 2007;14(4):682–92. 10.1038/sj.cdd.4402048

21. Anand R, Wai T, Baker MJ, Kladt N, Schauss AC, Rugarli E, et al. The i-AAA protease YME1L and OMA1 cleave OPA1 to balance mitochondrial fusion and fission. J Cell Biol. 2014;204(6):919–29. 10.1083/jcb.201308006

22. Baker MJ, Lampe PA, Stojanovski D, Korwitz A, Anand R, Tatsuta T, et al. Stress-induced OMA1 activation and autocatalytic turnover regulate OPA1-dependent mitochondrial dynamics. EMBO J. 2014;33(6):578–93. 10.1002/embj.201386474

23. Tang J, Zhang K, Dong J, Yan C, Hu C, Ji H, et al. Sam50-Mic19-Mic60 axis determines mitochondrial cristae architecture by mediating mitochondrial outer and inner membrane contact. Cell Death Differ. 2020;27(1):146–60. 10.1038/s41418-019-0345-2

24. Cho KF, Branon TC, Udeshi ND, Myers SA, Carr SA, Ting AY. Proximity labeling in mammalian cells with TurboID and split-TurboID. Nat Protoc. 2020;15(12):3971–99. 10.1038/s41596-020-0399-0

25. Lazarou M, Stojanovski D, Frazier AE, Kotevski A, Dewson G, Craigen WJ, et al. Inhibition of Bak activation by VDAC2 is dependent on the Bak transmembrane anchor. J Biol Chem. 2010;285(47):36876–83. 10.1074/jbc.M110.159301

26. Saunders TL, Windley SP, Gervinskas G, Balka KR, Rowe C, Lane R, et al. Exposure of the inner mitochondrial membrane triggers apoptotic mitophagy. Cell Death Differ. 2024;31(3):335–47. 10.1038/s41418-024-01260-2

27. Friedman JR, Lackner LL, West M, DiBenedetto JR, Nunnari J, Voeltz GK. ER tubules mark sites of mitochondrial division. Science. 2011;334(6054):358–62. 10.1126/science.1207385

28. De Vos K, Goossens V, Boone E, Vercammen D, Vancompernolle K, Vandenabeele P, et al. The 55-kDa Tumor Necrosis Factor Receptor Induces Clustering of Mitochondria through Its Membrane-proximal Region*. J Biol Chem. 1998;273(16):9673–80. 10.1074/jbc.273.16.9673

29. Hohorst L, Ros U, Garcia-Saez AJ. Mitochondrial dynamics and pore formation in regulated cell death pathways. Trends Biochem Sci. 2025;50(11):1001–14. 10.1016/j.tibs.2025.09.001

30. Brooks C, Wei Q, Feng L, Dong G, Tao Y, Mei L, et al. Bak regulates mitochondrial morphology and pathology during apoptosis by interacting with mitofusins. Proc Natl Acad Sci U S A. 2007;104(28):11649–54. 10.1073/pnas.0703976104

31. Rocha AL, Pai V, Perkins G, Chang T, Ma J, De Souza EV, et al. An Inner Mitochondrial Membrane Microprotein from the SLC35A4 Upstream ORF Regulates Cellular Metabolism. Journal of Molecular Biology. 2024;436(10):168559. 10.1016/j.jmb.2024.168559

32. Rocha AL, Schmedt C, Perkins G, Pinto A, Diedrich JK, Shan H, et al. Abnormal mitochondrial structure and function in brown adipose tissue of SLC35A4-MP knockout mice. Sci Adv. 2025;11(35):eads7381. doi:10.1126/sciadv.ads7381

33. Liu X, Kim CN, Yang J, Jemmerson R, Wang X. Induction of Apoptotic Program in Cell-Free Extracts: Requirement for dATP and Cytochrome c. Cell. 1996;86(1):147–57. 10.1016/S0092-8674(00)80085-9

34. Suzuki Y, Imai Y, Nakayama H, Takahashi K, Takio K, Takahashi R. A serine protease, HtrA2, is released from the mitochondria and interacts with XIAP, inducing cell death. Mol Cell. 2001;8(3):613–21. 10.1016/s1097-2765(01)00341-0

35. Adrain C, Creagh EM, Martin SJ. Apoptosis-associated release of Smac/DIABLO from mitochondria requires active caspases and is blocked by Bcl-2. EMBO J. 2001;20(23):6627–36.10.1093/emboj/20.23.6627

36. Tábara LC, Burr SP, Frison M, Chowdhury SR, Paupe V, Nie Y, et al. MTFP1 controls mitochondrial fusion to regulate inner membrane quality control and maintain mtDNA levels. Cell. 2024;187(14):3619–37.e27. 10.1016/j.cell.2024.05.017

37. Kraus F, Roy K, Pucadyil TJ, Ryan MT. Function and regulation of the divisome for mitochondrial fission. Nature. 2021;590(7844):57–66. 10.1038/s41586-021-03214-x

38. Adebayo M, Singh S, Singh AP, Dasgupta S. Mitochondrial fusion and fission: The fine-tune balance for cellular homeostasis. FASEB J. 2021;35(6):e21620. 10.1096/fj.202100067R

39. Wasiak S, Zunino R, McBride HM. Bax/Bak promote sumoylation of DRP1 and its stable association with mitochondria during apoptotic cell death. J Cell Biol. 2007;177(3):439– 50. 10.1083/jcb.200610042

40. Karbowski M, Lee Y-J, Gaume B, Jeong S-Y, Frank S, Nechushtan A, et al. Spatial and temporal association of Bax with mitochondrial fission sites, Drp1, and Mfn2 during apoptosis. J Cell Biol. 2002;159(6):931–8. 10.1083/jcb.200209124

41. Montessuit S, Somasekharan SP, Terrones O, Lucken-Ardjomande S, Herzig S, Schwarzenbacher R, et al. Membrane Remodeling Induced by the Dynamin-Related Protein Drp1 Stimulates Bax Oligomerization. Cell. 2010;142(6):889–901. 10.1016/j.cell.2010.08.017

42. He B, Yu H, Liu S, Wan H, Fu S, Liu S, et al. Mitochondrial cristae architecture protects against mtDNA release and inflammation. Cell Rep. 2022;41(10). 10.1016/j.celrep.2022.111774

43. Hassel KR, Brito-Estrada O, Makarewich CA. Microproteins: Overlooked regulators of physiology and disease. iScience. 2023;26(6):106781. 10.1016/j.isci.2023.106781

44. Anderson Douglas M, Anderson Kelly M, Chang C-L, Makarewich Catherine A, Nelson Benjamin R, McAnally John R, et al. A Micropeptide Encoded by a Putative Long Noncoding RNA Regulates Muscle Performance. Cell. 2015;160(4):595–606. 10.1016/j.cell.2015.01.009

45. Anderson DM, Makarewich CA, Anderson KM, Shelton JM, Bezprozvannaya S, Bassel-Duby R, et al. Widespread control of calcium signaling by a family of SERCA-inhibiting micropeptides. Sci Signaling. 2016;9(457):ra119–ra. doi:10.1126/scisignal.aaj1460

46. Liang C, Zhang S, Robinson D, Ploeg MV, Wilson R, Nah J, et al. Mitochondrial microproteins link metabolic cues to respiratory chain biogenesis. Cell Rep. 2022;40(7). 10.1016/j.celrep.2022.111204

47. Dennerlein S, Poerschke S, Oeljeklaus S, Wang C, Richter-Dennerlein R, Sattmann J, et al. Defining the interactome of the human mitochondrial ribosome identifies SMIM4 and TMEM223 as respiratory chain assembly factors. Elife. 2021;10. 10.7554/eLife.68213

48. Rathore A, Chu Q, Tan D, Martinez TF, Donaldson CJ, Diedrich JK, et al. MIEF1 Microprotein Regulates Mitochondrial Translation. Biochem. 2018;57(38):5564–75. 10.1021/acs.biochem.8b00726

49. Dibley MG, Formosa LE, Lyu B, Reljic B, McGann D, Muellner-Wong L, et al. The Mitochondrial Acyl-carrier Protein Interaction Network Highlights Important Roles for LYRM Family Members in Complex I and Mitoribosome Assembly. Mol Cell Proteomics. 2020;19(1):65–77. 10.1074/mcp.RA119.001784

50. Olichon A, Baricault L, Gas N, Guillou E, Valette A, Belenguer P, et al. Loss of OPA1 perturbates the mitochondrial inner membrane structure and integrity, leading to cytochrome c release and apoptosis. J Biol Chem. 2003;278(10):7743–6. 10.1074/jbc.C200677200

51. Song Z, Chen H, Fiket M, Alexander C, Chan DC. OPA1 processing controls mitochondrial fusion and is regulated by mRNA splicing, membrane potential, and Yme1L. J Cell Biol. 2007;178(5):749–55. 10.1083/jcb.200704110

52. Cogliati S, Frezza C, Soriano ME, Varanita T, Quintana-Cabrera R, Corrado M, et al. Mitochondrial cristae shape determines respiratory chain supercomplexes assembly and respiratory efficiency. Cell. 2013;155(1):160–71. 10.1016/j.cell.2013.08.032

53. Del Dotto V, Mishra P, Vidoni S, Fogazza M, Maresca A, Caporali L, et al. OPA1 Isoforms in the Hierarchical Organization of Mitochondrial Functions. Cell Rep. 2017;19(12):2557–71. 10.1016/j.celrep.2017.05.073

54. Ding C, Wu Z, Huang L, Wang Y, Xue J, Chen S, et al. Mitofilin and CHCHD6 physically interact with Sam50 to sustain cristae structure. Sci Rep. 2015;5:16064. 10.1038/srep16064

55. Cipolat S, Rudka T, Hartmann D, Costa V, Serneels L, Craessaerts K, et al. Mitochondrial Rhomboid PARL Regulates Cytochrome c Release during Apoptosis via OPA1-Dependent Cristae Remodeling. Cell. 2006;126(1):163–75. 10.1016/j.cell.2006.06.021

56. Pereira RO, Olvera AC, Marti A, Fang S, White JR, Westphal M, et al. OPA1 Regulates Lipid Metabolism and Cold-Induced Browning of White Adipose Tissue in Mice. Diabetes. 2022;71(12):2572–83. 10.2337/db22-0450

57. Liu GY, Moon SH, Jenkins CM, Li M, Sims HF, Guan S, et al. The phospholipase iPLA(2)γ is a major mediator releasing oxidized aliphatic chains from cardiolipin, integrating mitochondrial bioenergetics and signaling. J Biol Chem. 2017;292(25):10672–84. 10.1074/jbc.M117.783068

58. He J, Mao CC, Reyes A, Sembongi H, Di Re M, Granycome C, et al. The AAA+ protein ATAD3 has displacement loop binding properties and is involved in mitochondrial nucleoid organization. J Cell Biol. 2007;176(2):141–6. 10.1083/jcb.200609158

59. Ishihara T, Ban-Ishihara R, Ota A, Ishihara N. Mitochondrial nucleoid trafficking regulated by the inner-membrane AAA-ATPase ATAD3A modulates respiratory complex formation. Proc Natl Acad Sci U S A. 2022;119(47):e2210730119. doi:10.1073/pnas.2210730119

60. Peralta S, Goffart S, Williams SL, Diaz F, Garcia S, Nissanka N, et al. ATAD3 controls mitochondrial cristae structure in mouse muscle, influencing mtDNA replication and cholesterol levels. J Cell Sci. 2018;131(13). 10.1242/jcs.217075

61. Sen A, Kallabis S, Gaedke F, Jüngst C, Boix J, Nüchel J, et al. Mitochondrial membrane proteins and VPS35 orchestrate selective removal of mtDNA. Nat Commun. 2022;13(1):6704. 10.1038/s41467-022-34205-9

62. Ran FA, Hsu PD, Wright J, Agarwala V, Scott DA, Zhang F. Genome engineering using the CRISPR-Cas9 system. Nat Protoc. 2013;8(11):2281–308. 10.1038/nprot.2013.143

63. Formosa LE, Maghool S, Sharpe AJ, Reljic B, Muellner-Wong L, Stroud DA, et al. Mitochondrial COA7 is a heme-binding protein with disulfide reductase activity, which acts in the early stages of complex IV assembly. Proc Natl Acad Sci U S A. 2022;119(9):e2110357119. doi:10.1073/pnas.2110357119

64. Labun K, Montague TG, Krause M, Torres Cleuren YN, Tjeldnes H, Valen E. CHOPCHOP v3: expanding the CRISPR web toolbox beyond genome editing. Nucleic Acids Res. 2019;47(W1):W171–W4. 10.1093/nar/gkz365

65. Formosa LE, Muellner-Wong L, Reljic B, Sharpe AJ, Jackson TD, Beilharz TH, et al. Dissecting the Roles of Mitochondrial Complex I Intermediate Assembly Complex Factors in the Biogenesis of Complex I. Cell Rep. 2020;31(3):107541. 10.1016/j.celrep.2020.107541

66. Johnston AJ, Hoogenraad J, Dougan DA, Truscott KN, Yano M, Mori M, et al. Insertion and assembly of human tom7 into the preprotein translocase complex of the outer mitochondrial membrane. J Biol Chem. 2002;277(44):42197–204. 10.1074/jbc.M205613200

67. Schägger H, von Jagow G. Tricine-sodium dodecyl sulfate-polyacrylamide gel electrophoresis for the separation of proteins in the range from 1 to 100 kDa. Anal Biochem. 1987;166(2):368–79. 10.1016/0003-2697(87)90587-2

68. Wittig I, Braun HP, Schägger H. Blue native PAGE. Nat Protoc. 2006;1(1):418–28. 10.1038/nprot.2006.62

69. McKenzie M, Lazarou M, Thorburn DR, Ryan MT. Analysis of mitochondrial subunit assembly into respiratory chain complexes using Blue Native polyacrylamide gel electrophoresis. Anal Biochem. 2007;364(2):128–37. 10.1016/j.ab.2007.02.022

70. Kamerkar SC, Kraus F, Sharpe AJ, Pucadyil TJ, Ryan MT. Dynamin-related protein 1 has membrane constricting and severing abilities sufficient for mitochondrial and peroxisomal fission. Nat Commun. 2018;9(1):5239. 10.1038/s41467-018-07543-w

71. Schindelin J, Arganda-Carreras I, Frise E, Kaynig V, Longair M, Pietzsch T, et al. Fiji: an open-source platform for biological-image analysis. Nat Methods. 2012;9(7):676–82. 10.1038/nmeth.2019

72. Grimm JB, Tkachuk AN, Patel R, Hennigan ST, Gutu A, Dong P, et al. Optimized red-absorbing dyes for imaging and sensing. J Am Chem Soc. 2023;145(42):23000–13.

73. Danne JC, Crawford SA, Templin R, Clark J, Oorschot V, Ramm G. Preparation of cells for transmission electron microscopy ultrastructural analysis V.1 protocolsio. 2022. 10.17504/protocols.io.j8nlk47ewg5r/v1

74. Rappsilber J, Ishihama Y, Mann M. Stop and Go Extraction Tips for Matrix-Assisted Laser Desorption/Ionization, Nanoelectrospray, and LC/MS Sample Pretreatment in Proteomics. Anal Chem. 2003;75(3):663–70. 10.1021/ac026117i

75. Szklarczyk D, Kirsch R, Koutrouli M, Nastou K, Mehryary F, Hachilif R, et al. The STRING database in 2023: protein-protein association networks and functional enrichment analyses for any sequenced genome of interest. Nucleic Acids Res. 2023;51(D1):D638–d46. 10.1093/nar/gkac1000

