## Supplementary Figures & Tables for "Proximity labelling of the BAK macropore uncovers a new role for SLC35A4-MP in mitochondrial dynamics"

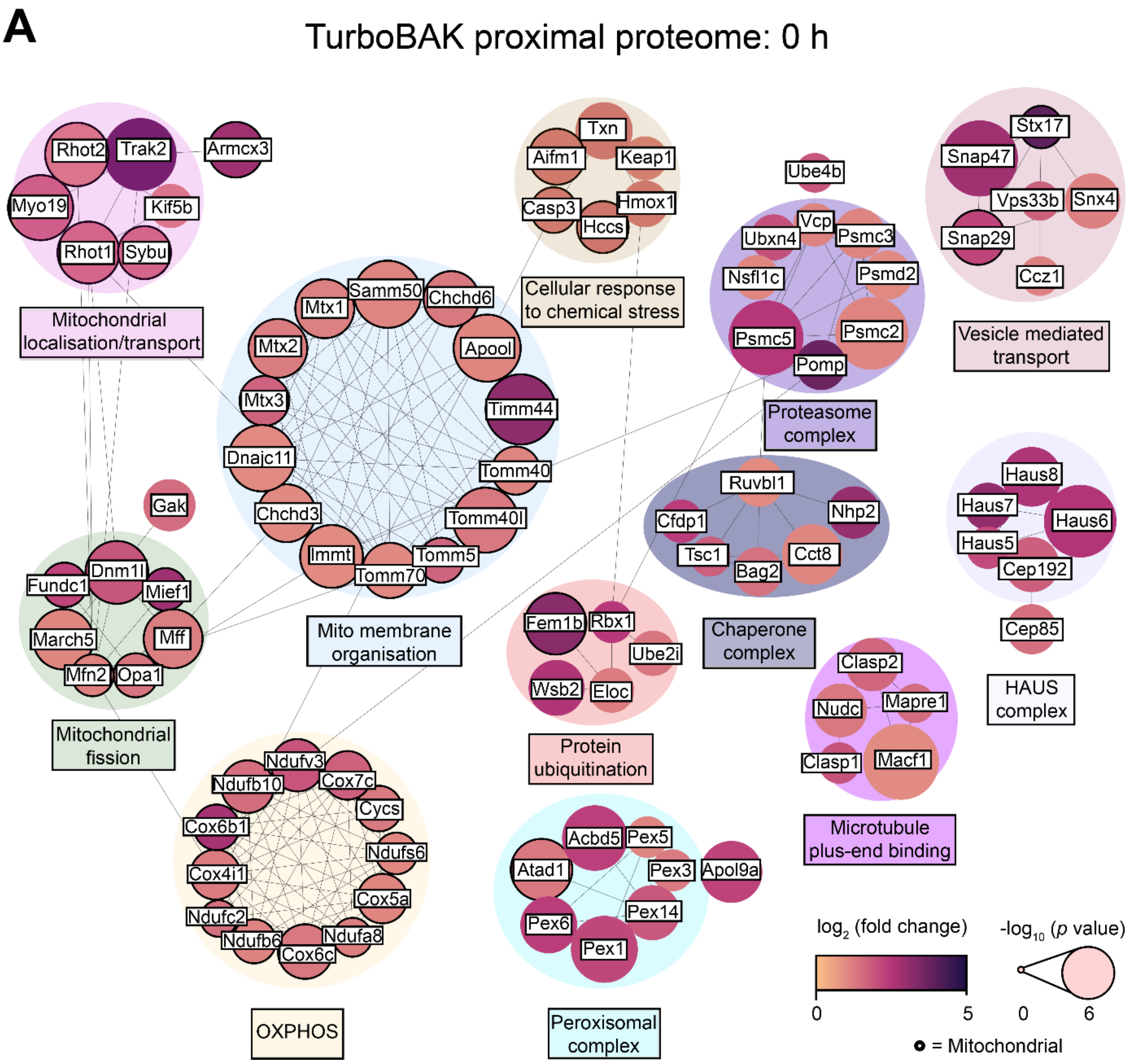


**Figure S1: A** Protein-protein interaction network of significantly enriched proteins at 0 h + biotin. Connections indicate high confidence interactions. Only protein clusters with ≥ 5 enriched proteins are shown. Clusters have been annotated with enriched GO terms. Cytoscape was used to colour and size the nodes based on the strength of enrichment ($\log_2$ (fold change) and $p$ value respectively). Known mitochondrial annotations are indicated.

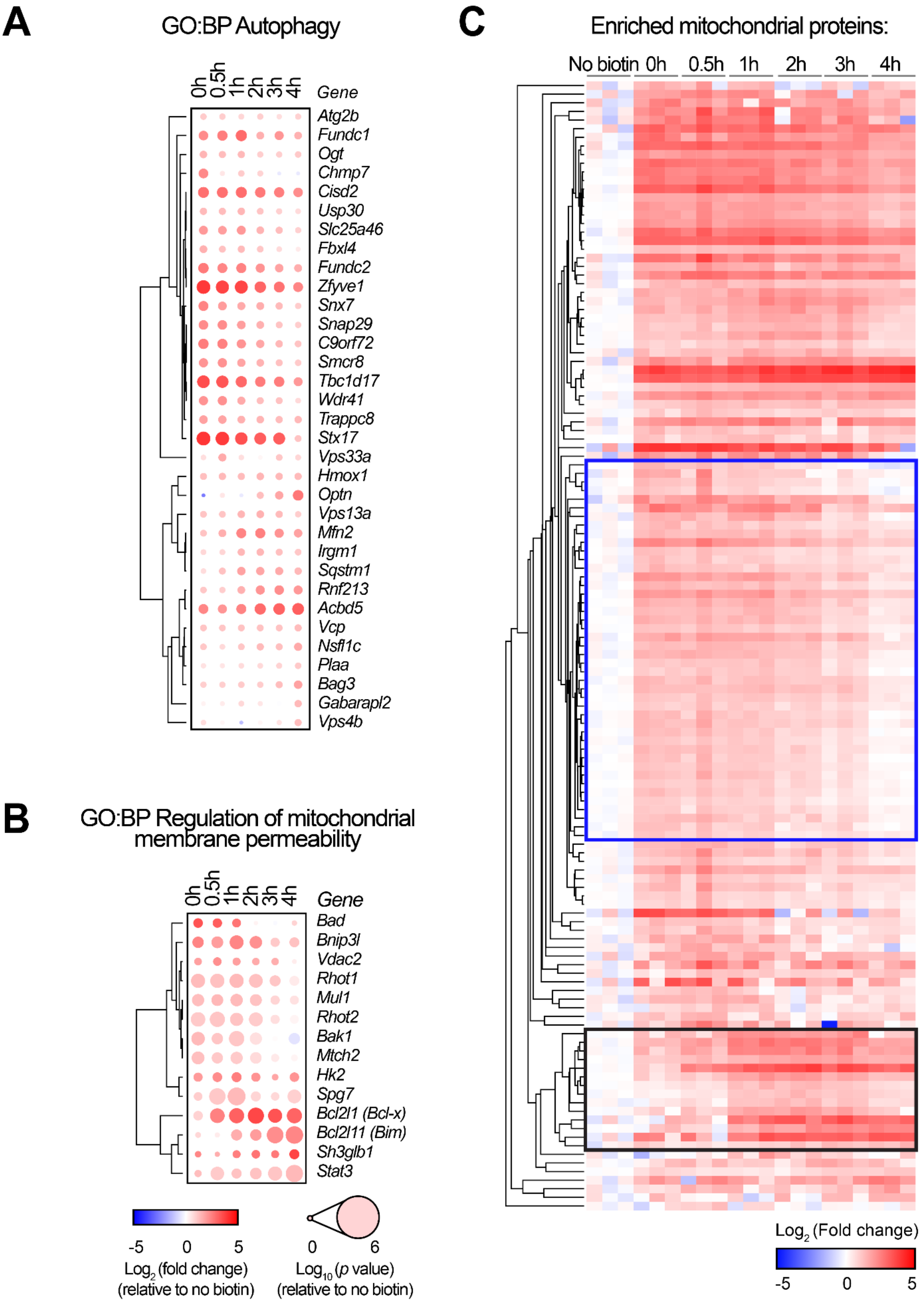
A
GO:BP Autophagy
0h
0.5h
1h
2h
3h
4h
Gene
Atg2b
Fundc1
Ogt
Chmp7
Cisd2
Usp30
Slc25a46
Fbxl4
Fundc2
Zfyve1
Snx7
Snap29
C9orf72
Smcr8
Tbc1d17
Wdr41
Trappc8
Stx17
Vps33a
Hmox1
Optn
Vps13a
Mfn2
Irgm1
Sqstm1
Rnf213
Acbd5
Vcp
Nsfl1c
Plaa
Bag3
Gabarapl2
Vps4b
B
GO:BP Regulation of mitochondrial membrane permeability
0h
0.5h
1h
2h
3h
4h
Gene
Bad
Bnip3l
Vdac2
Rhot1
Mul1
Rhot2
Bak1
Mtch2
Hk2
Spg7
Bcl2l1 (Bcl-x)
Bcl2l11 (Bim)
Sh3glb1
Stat3
-5 0 5
Log2 (fold change) (relative to no biotin)
0 6
Log10 (p value) (relative to no biotin)
C
Enriched mitochondrial proteins:
No biotin
0h
0.5h
1h
2h
3h
4h
Log2 (Fold change)
-5 0 5

**Figure S2: A-B.** Significantly enriched proteins with the GO Biological Process term **A)** **"**Autophagy" or **B)** **"**Regulation of mitochondrial membrane permeability". Colour and size indicate the strength of enrichment ($log_2$ (fold change) and *p* value respectively. Proteins were hierarchically clustered using one minus Pearson correlation. **C.** Hierarchical clustering (one minus Pearson correlation) of mitochondrial proteins significantly enriched at any timepoint. The cluster of decreased mitochondrial proteins is highlighted in blue, and the cluster of increased mitochondrial proteins are highlighted in black.

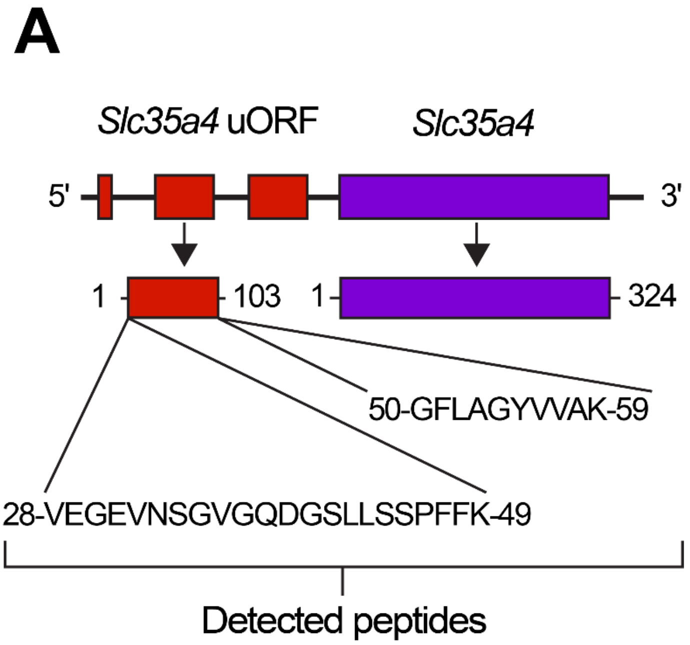


**Figure S3A.** Schematic of the *Slc35a4* gene locus. Detected SLC35A4 peptides are annotated.

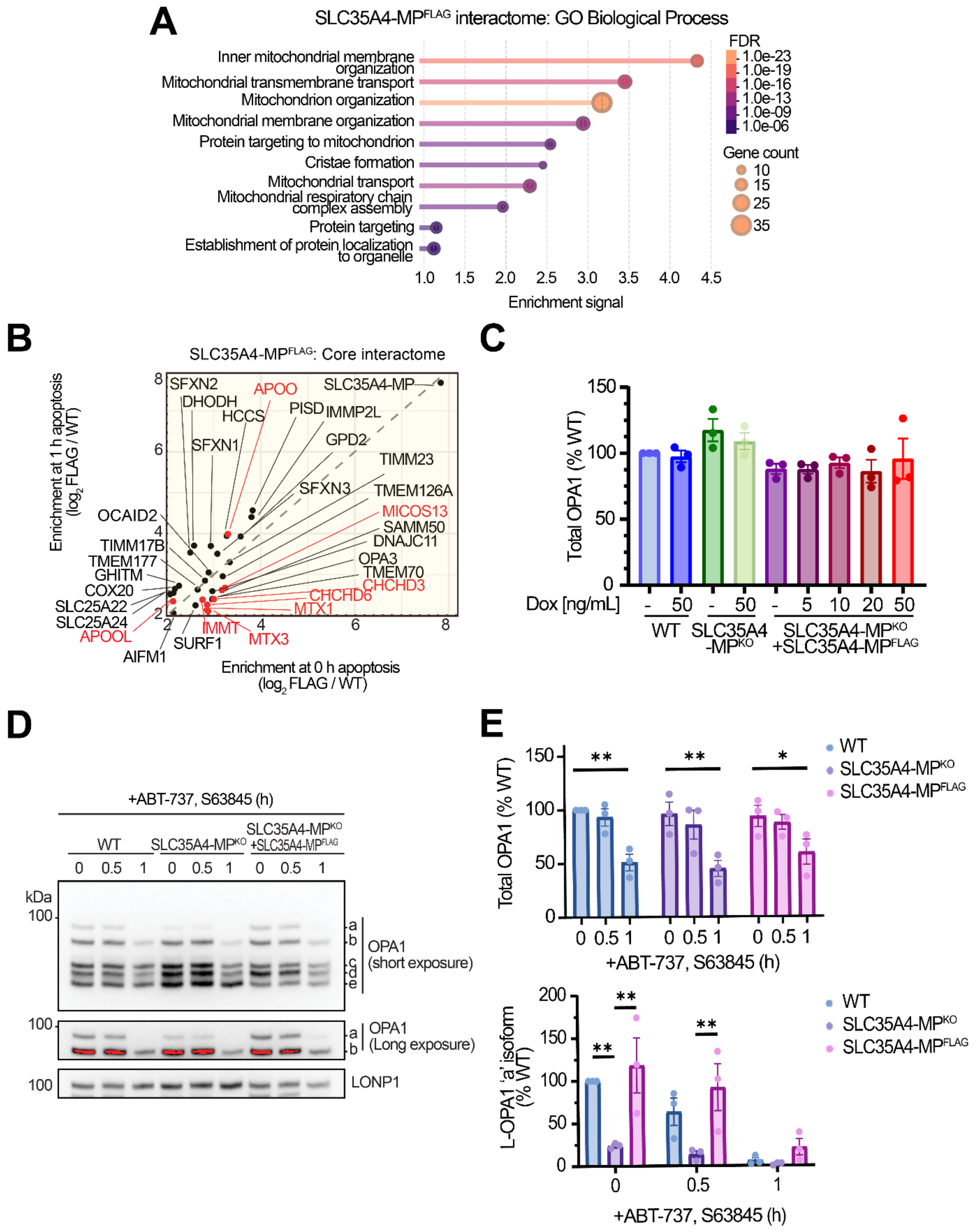
A
SLC35A4-MPFLAG interactome: GO Biological Process
Inner mitochondrial membrane organization
Mitochondrial transmembrane transport
Mitochondrion organization
Mitochondrial membrane organization
Protein targeting to mitochondrion
Cristae formation
Mitochondrial transport
Mitochondrial respiratory chain complex assembly
Protein targeting
Establishment of protein localization to organelle
Enrichment signal
FDR
Gene count
B
SLC35A4-MPFLAG: Core interactome
Enrichment at 1 h apoptosis (log2 FLAG / WT)
Enrichment at 0 h apoptosis (log2 FLAG / WT)
SFXN2
APOO
SLC35A4-MP
DHODH
HCCS
PISD
IMMP2L
SFXN1
GPD2
TIMM23
SFXN3
TMEM126A
OCAID2
MICOS13
SAMM50
DNAJC11
TIMM17B
TMEM177
GHITM
COX20
SLC25A22
SLC25A24
APOOL
AIFM1
SURF1
IMMT
MTX3
MTX1
CHCHD6
CHCHD3
TMEM70
OPA3
C
Total OPA1 (% WT)
Dox [ng/mL]
WT
SLC35A4-MPKO
SLC35A4-MPKO +SLC35A4-MPFLAG
D
+ABT-737, S63845 (h)
WT
SLC35A4-MPKO
SLC35A4-MPKO +SLC35A4-MPFLAG
kDa
OPA1 (short exposure)
OPA1 (Long exposure)
LONP1
E
Total OPA1 (% WT)
WT
SLC35A4-MPKO
SLC35A4-MPFLAG
+ABT-737, S63845 (h)
L-OPA1 'a' isoform (% WT)

**Figure S4A.** GO Biological Process enrichment analysis of SLC35A4-MP$^{FLAG}$ interactome at steady-state. **B.** The Core SLC35A4-MP$^{FLAG}$ interactome (yellow region from Fig. 4B). Comparison of $\log_2$ (Flag / WT) enrichment of the SLC35A4-MP$^{FLAG}$ interactome at 0 h and 1 h apoptosis. Enriched components of the MICOS-MIB complex are highlighted in red. **C.** Quantification of total OPA1 abundance following induction of SLC35A4-MP$^{FLAG}$ expression at increasing concentrations of doxycycline **D.** SDS-PAGE and immunoblotting of isolated mitochondria from WT, SLC35A4–MP$^{KO}$ and SLC35A4-MP rescue (SLC35A4-MP$^{FLAG}$; 5 ng / ml doxycycline for 48 h) U2OS cells, treated with ABT737 [10 μM], S63845 [2 μM] and QVD-OPh [20 μM] (1 h pre-treat). Red indicates saturated pixels. LONP1 serves as the loading control. Representative of n = 3 independent experiments. **E.** Densitometric quantification of L-OPA1 'a' isoform or total OPA1. Pixel intensity was normalised to LONP1 levels and displayed relative to the 0 h WT control. * = $p < 0.05$, ** = $p < 0.01$ by one-way ANOVA with Tukey's multiple comparison test. Quantified from n = 3 independent experiments.

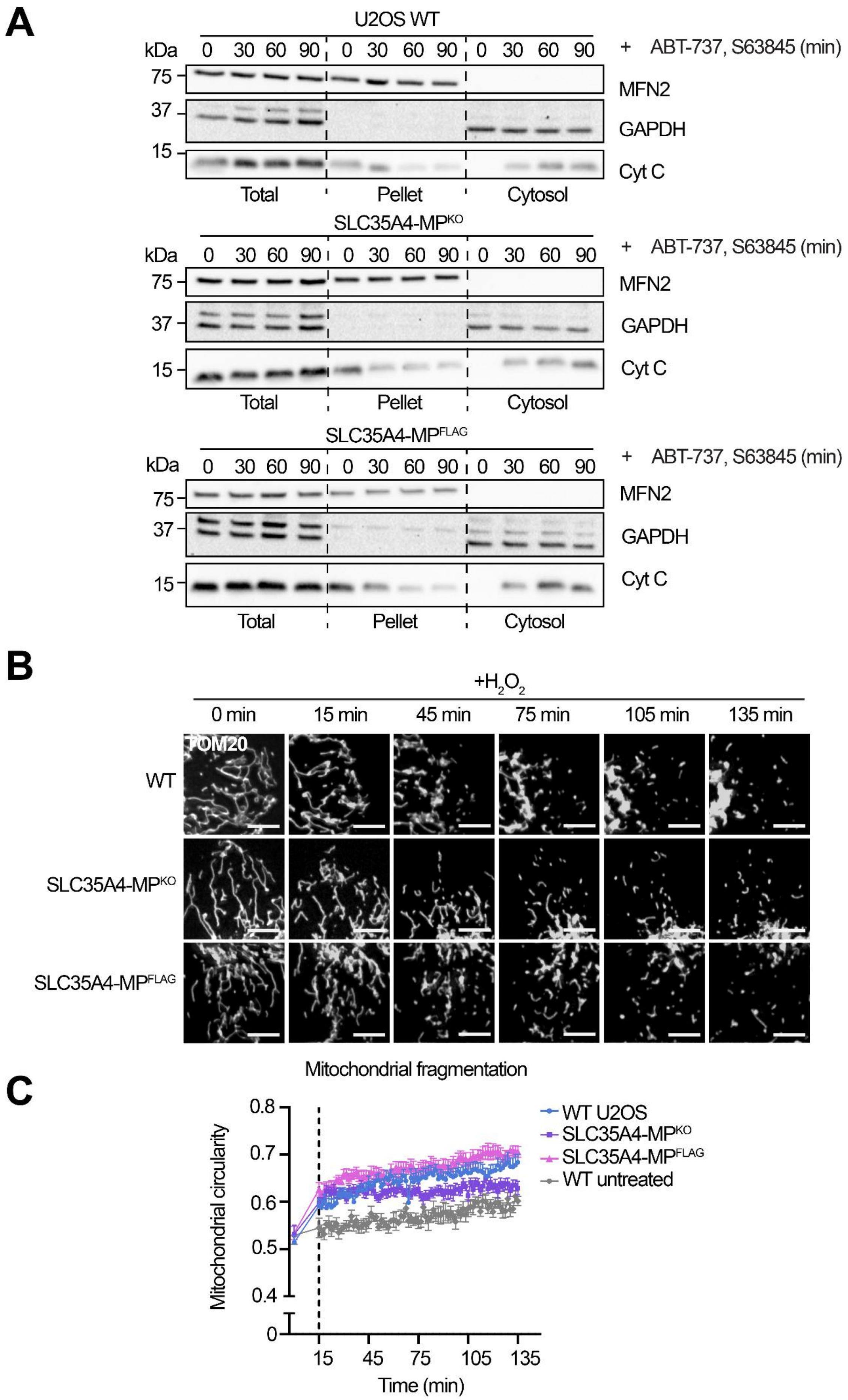
A
U2OS WT
kDa
0 30 60 90 0 30 60 90 0 30 60 90
+ ABT-737, S63845 (min)
75
37
15
MFN2
GAPDH
Cyt C
Total
Pellet
Cytosol
SLC35A4-MP^KO
SLC35A4-MP^FLAG
B
+H2O2
0 min
15 min
45 min
75 min
105 min
135 min
TOM20
WT
SLC35A4-MP^KO
SLC35A4-MP^FLAG
C
Mitochondrial fragmentation
Mitochondrial circularity
0.8
0.7
0.6
0.5
0.4
0
15
45
75
105
135
Time (min)
WT U2OS
SLC35A4-MP^KO
SLC35A4-MP^FLAG
WT untreated

**Supplementary figure 5A.** SDS-PAGE separation and immunoblotting of sub-cell fractionated U2OS WT, SLC35A4-MP$^{KO}$ and SLC35A4-MP rescue cells expressing endogenous levels of SLC35A4-MP$^{FLAG}$ (5 ng / ml doxycycline for 48 h), treated with ABT737 [10 µM], S63845 [2 µM] and QVD-OPh [20 µM] (1 h pre-treat). Representative of n = 2 independent experiments. **B.** Live cell imaging of U2OS WT, SLC35A4-MP$^{KO}$ or SLC35A4-MP rescue (SLC35A4-MP$^{FLAG}$; 5 ng / ml doxycycline for 48 h) stably expressing TOM20$^{Halo}$ following treatment with $H_2O_2$ {500 µM]. Representative images of n = 3 independent experiments. Scale bars indicate 10 µm. **C.** Quantification of mitochondrial circularity, denoting $H_2O_2$ induced mitochondrial network collapse. Quantification was performed on 10-15 cells from n = 3 independent experiments. Data points indicate mean ± SEM.

| **Antibodies: Immunoblots (Dilution)** | **Source** | **Identifier** |
|---|---|---|
| Anti-ATP5A (1:1000) | Abcam | Ab14748 |
| Anti-BAK (1:1000) | Made in house | NA |
| Anti-Cytochrome *c* (1:300) | Invitrogen | 45-6100 |
| Anti-FLAG (1:1000) | Sigma-Aldrich | F1804 |
| Anti-GAPDH (1:2000) | Cell Signalling | 2118S |
| Anti-LONP1 (1:1000) | Cell Signalling | 28020T |
| Anti-MFN2 (1:1000) | Cell Signalling | 9482 |
| Anti-MIC10 (1:2000) | Aviva Systems Biology | ARP44801_P050 |
| Anti-MIC19 (1:1000) | Abclonal | A8584 |
| Anti-MIC27 (1:500) | Santa Cruz Biotechnology | sc-390958 |
| Anti-MIC60 (1:1000) | Abcam | Ab48139 |
| Anti-NDUFA9 (1:500) | Made in house | NA |
| Anti-OPA1 (1:1000) | BD Biosciences | 612606 |
| Anti-SAM50 (1:500) | Made in house | NA |
| Anti-SDHA (1:1000) | Abcam | Ab14715 |
| Anti-SLC35A4-MP (1:1000) | Made in house | NA |
| **Antibodies: Immunofluorescence assay (Dilution)** | | |
| Anti-DNA (1:200) | ThermoFisher Scientific | 61014PROGEN |
| Anti-HA (1:250) | Cell Signalling | 2367S |
| Anti-FLAG (1:500) | Sigma-Aldrich | F1804 |
| Anti-TOM20 (1:500) | Santa Cruz Biotechnology | sc-11415 |

**Supplementary table 1.** Primary antibodies used for immunoblotting and immunofluorescence assays.
